# Coenzyme A depletion causes antibiotic tolerance in *Pseudomonas aeruginosa*

**DOI:** 10.1101/2025.10.24.684135

**Authors:** Pablo Manfredi, Isabella Santi, Enea Maffei, Hector Arturo Hernandez Gonzalez, Shannon Conroy, Emanuelle Lezan, Erik Ahrnè, Nicole Thürkauf, Simon van Vliet, Nicola Zamboni, Alexander Schmidt, Urs Jenal

## Abstract

The widespread use of antibiotics promotes both resistance and tolerance. While resistance enables bacterial growth in the presence of drugs, tolerance allows survival during treatment, generating persisters that seed relapse and promote resistance. Despite its clinical relevance, the molecular basis of tolerance remains poorly understood. Using proteomic and metabolomic profiling combined with machine learning, we identified thiol oxidation as a robust predictor of tolerance in the human pathogen *Pseudomonas aeruginosa*. Single-cell analyses established a direct link between thiol oxidation and drug survival, indicating that redox imbalance drives persistence. Whereas depletion of coenzyme A (CoA), a central thiol-containing metabolite, scaled with tolerance, restoring CoA using engineered catalysts from *Staphylococcus aureus* abolished tolerance, establishing a causal relation between CoA availability and drug susceptibility. Thiol-based predictors also accurately capture tolerance of clinical *P. aeruginosa* isolates. These findings establish CoA-centered redox control as a key determinant of tolerance, opening opportunities for diagnostics and therapeutic interventions to prevent infection relapses.

## Introduction

The widespread use of antibiotics promotes the evolution and dissemination of drug-specific resistance mechanisms. In contrast, tolerance generally protects against different classes of bactericidal antibiotics and provides a particularly strong selective advantage during combination therapy or sequential drug application ^1,2^. Bacterial cultures frequently exhibit subpopulations of highly tolerant persisters, which can initiate post-treatment regrowth and promote the emergence of resistance ^2–4^. Variants with increased tolerance rapidly emerge during recurrent exposure of bacterial cultures to high concentrations of antibiotics. This process precedes and promotes the fixation of resistance-conferring mutations and is often followed by reversion of the fitness-costly tolerance phenotype ^2,4^. Nevertheless, resistance and tolerance can still frequently coexist ^2^. Accordingly, tolerance is experimentally defined as increased drug survival in strains that share identical minimal inhibitory concentrations (MICs).

Although multi-drug tolerance is well described in a range of human pathogens ^2,5–7^, little is known about the physiological mechanisms responsible for this phenomenon and the respective molecular drivers have remained controversial ^8–10^. Proposed tolerance drivers include toxin–antitoxin systems ^4,11,12^, the SOS-response ^13^, the alarmone (p)ppGpp ^14^, changes in intracellular pH homeostasis ^15^, as well as fluctuating levels of ATP and energy homeostasis ^16^. Moreover, stochastic cell-to-cell variations of drug targets ^17^, various forms of stress associated with host environments ^18^, or random errors in cell replication ^19^ were proposed to cause drug tolerance in bacteria. Sub-lethal stressors and the resulting cellular responses were also proposed to lower antimicrobial efficiency, but this relationship remained unclear as cells primed with acid, temperature or osmotic shock displayed both increased or decreased survival to antibiotics ^20–23^. More specific forms of stress associated with antibiotic tolerance include redox imbalances caused by metabolic alterations ^24^ or by the host’s immune response ^25^. Intriguingly, redox stress and reactive oxygen species were also proposed to promote cell killing by bactericidal antibiotics ^26,27^, arguing that bacteria can either be protected or killed depending on the context and the degree of the imposed redox stress.

To better define the physiological mechanisms underlying drug tolerance, we employed a non-biased approach to study the physiological characteristics of a set of persister strains with varying levels of drug tolerance from the high priority pathogen *Pseudomonas aeruginosa* ^28^. By applying machine learning algorithms to select omics-derived variables predictive of drug tolerance, we identified a robust correlation between antimicrobial tolerance and the degree of thiol oxidation. This relationship was evident not only in tolerance levels across different mutants but also within clonal populations where tolerant subpopulations consistently exhibited elevated thiol oxidation states. Notably, we found that the levels of coenzyme A (CoA), a central thiol-containing cellular metabolite, closely correlate with the degree of *P. aeruginosa* drug tolerance. By expressing heterologous enzymes that specifically restore CoA availability in strains with high tolerance, we were able to demonstrate a direct causal relationship between CoA levels and drug sensitivity. We show that these findings are applicable to a broad range of clinical *P. aeruginosa* isolates with increased tolerance, providing a foundation for developing new diagnostic tools and treatments aimed at preventing infection relapses and the emergence of antimicrobial resistance.

## Results

### Identification of hyper-tolerant *P. aeruginosa* lineages before treatment with antibiotics

To investigate the molecular basis of antibiotic tolerance, we made use of hyper-tolerant isolates that were obtained by recurrent exposure of *P. aeruginosa* to high concentrations of different antibiotics (Figure 1a) ^2,29^. Mutations conferring drug tolerance (henceforth referred as * alleles) all mapped to proteins involved in redox reactions, respiration or energy production ^2^. This includes mutations in NuoN, a transmembrane subunit of the respiratory complex I; CcmG and CcmF, enzymes of the cytochrome c maturation pathway; NatT, a RES domain toxin involved in NAD^+^ and NADP^+^ degradation ^11^ and CoaD, a feedback-regulated enzyme of coenzyme A (CoA) biosynthesis ^30^(Figure 1b). The only exception to this redox-associated pattern were mutations in ParS, a sensor histidine kinase regulating outer membrane permeability and expression of efflux pumps ^31^. These mutations conferred different levels of tolerance across different classes of antibiotics upon entry in stationary phase (Figures 1c,d, S1a) without causing increased resistance (Figure S1b).

**Figure 1:**
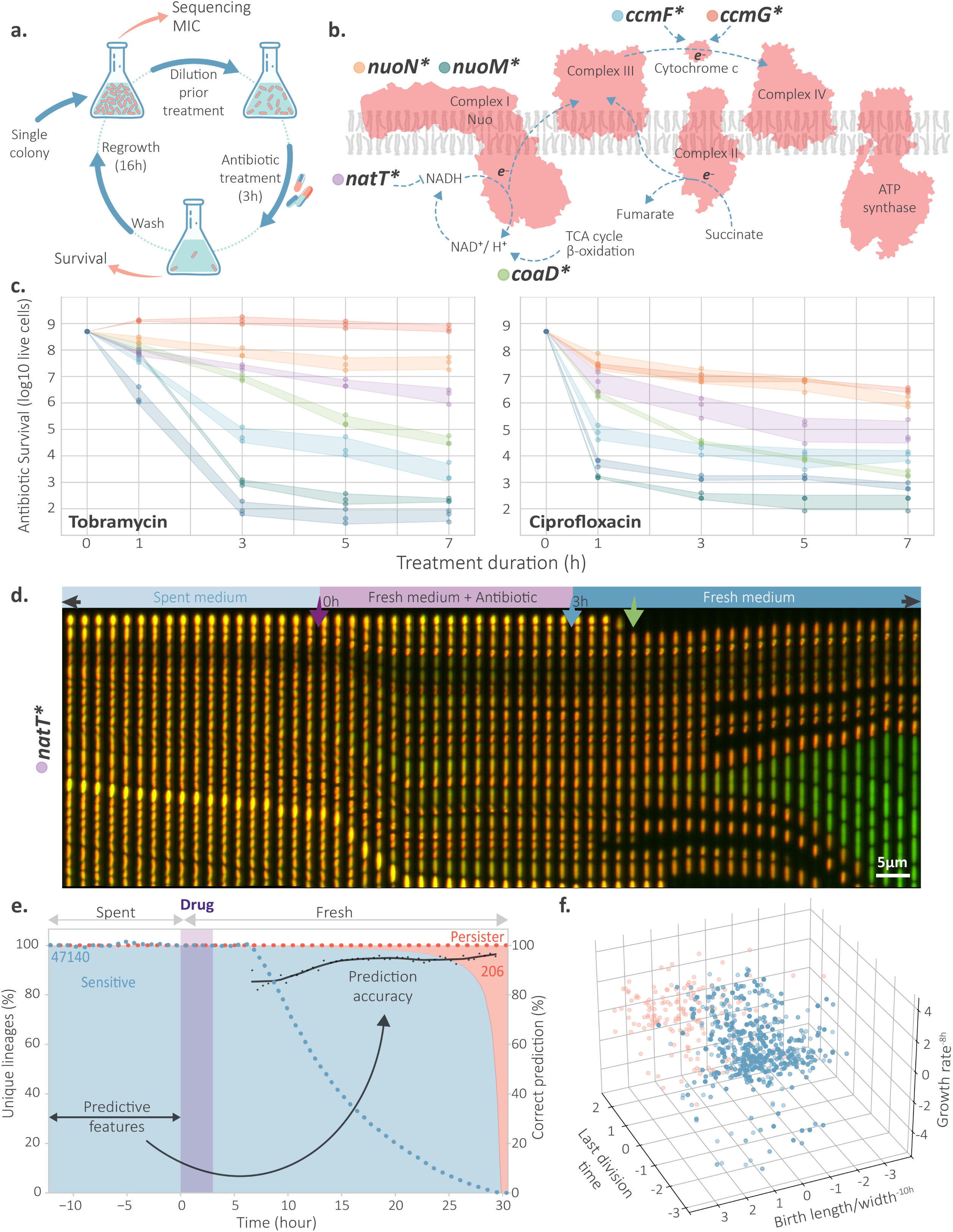
Mutations conferring antimicrobial tolerance map to the respiratory chain of *P. aeruginosa*. **(a)** Schematic of experimental evolution of *P. aeruginosa* tolerance with periodic exposures to bactericidal antibiotics ^2^. **(b)** Mutations causing hyper-tolerance map to components of the respiratory chain. Protein complexes of the electron transport chain (red) and electron fluxes (blue arrows) are shown with hyper-tolerance mutations indicated with colored dots. **(c)** Killing kinetics of the different *P. aeruginosa* hyper-tolerant strains upon exposure to tobramycin (16 μg/ml) (left) or ciprofloxacin (2.5 μg/ml) (right). Colored dots (n=3), lines (LOESS regressions) and areas (95% confidence intervals) refer to mutant strains shown in panel (b) with the same color code. **(d)** Time-lapse microscopy of a microchannel exhibiting a persister cell from the *natT*\* hyper-tolerant lineage expressing the growth rate reporter TIMER ^32^. The Kymograph shows images taken every 10 minutes for 10 hours before and after antibiotic treatment. A phases of microchannels bacterial seeding with fresh medium (3 h, not shown) was followed by a stepwise increase of 100% spent medium (12.5 h), exposure to fresh medium containing tobramycin (16 μg/ml) for 3 h, and addition of fresh medium without antibiotics. Actively growing cells show green TIMER fluorescence, while cells with reduced growth rates are yellow or red. **(e)** Quantification of single cell tracking of susceptible (blue, 47140 cells) and drug tolerant (red, 206 cells) cells before, during and after treatment as shown in (d). Areas indicate the fraction of susceptible (blue) and drug tolerant cells (red) at different time points of the experiment. Dotted lines show the fraction of tolerant (red) and susceptible cells (blue) that were still visible at a given time point during treatment. The treatment window is indicated by a purple box. Microscopic features from the pre-treatment phase are exploited by the plsr approach to predict persister lineages. Prediction accuracy of the plsr model (right Y-axis) for survival and killing is shown as black dots for sensitive cells disintegrating at different time points (black line, LOESS regression). **(f)** Plot of the most distinguishable individual lineages classified as susceptible (blue dots) and persister (pink dots), based on the three most discriminating features identified by the PLSR models predicting survival following treatment-mediated killing : i) normalized ratio between cell size at birth divided by cell width 10 h prior treatment (X-axis); ii) normalized growth rate 8 h prior treatment (Y-axis); time of last division prior treatment with higher values being closer to treatment window (Z-axis).

We first used microfluidics to address if we can distinguish hyper-tolerant *P. aeruginosa* from their susceptible counterparts without selection by killing. An important experimental challenge when investigating hyper-tolerant strains is their genetic instability, which leads to the rapid invasion of compensatory mutations in the absence of selection. For example, without periodic antibiotic exposure, a second site mutation in *nuoM* invades the *nuoN\** hyper-tolerant strain of *P. aeruginosa*, alleviating its fitness costs but also largely abolishing its overall tolerance (Figure 1c) ^2^. Because of this genetic instability, experiments that required non-selective conditions focused on more stable hyper-tolerance alleles like *natT\** (Figure 1c) ^11^.

To visualize the formation of hyper-tolerant persisters at the single cell level in a microchannel “mother machine” fluidic device, we engineered a derivative of the *natT\** mutant lacking adhesive properties and motility (Δ*pelA-*Δ*pslABC-*Δ*pilA-*Δ*fliC*) and constitutively expressing the fluorescent growth indicator TIMER (*CTX::TIMER-DsRed*) ^32^. This setup allows analyzing 10’000s of cells in parallel, tracking rare persister lineages before, during and after drug treatment ^11^. While persisters were never observed in *P. aeruginosa* wild type, analysis of 47’140 lineages identified a total of 206 persisters in the *natT\** mutant, closely aligning with persister frequencies observed in bulk populations (Figure 1c,d,e). To investigate if persisters can be identified prior to treatment onset, we applied a simple machine learning approach and models trained by time-resolved microscopy data to predict persister fate based on pre-treatment lineage-derived features (see Materials and Methods). The models demonstrated robust predictive performance, achieving recall rates between 80% and 96% (Figure 1e). While several highly predictive features were associated with dynamic changes of cell length during the transition into stationary phase, the most robust predictor of persistence was the time of the last cell division. Persister cells tended to undergo their final division significantly closer to the onset of antibiotic treatment, suggesting that they maintained active cell division for a prolonged period during nutrient starvation (Figure 1f, S1c,d,e). Importantly, these experiments demonstrate that persisters can be identified *before* bacteria are being treated with antibiotics, ruling out that their appearance, frequency or physiology is being skewed by post-treatment effects ^33^.

### A unique proteomic signature quantitatively predicts *P. aeruginosa* antibiotic tolerance

Because alleles that enhance survival cluster around components of the *P. aeruginosa* electron transport chain (Figure 1b), we hypothesized that tolerance may arise from shared physiological properties. To test this, we employed a non-biased, proteomics-based regression approach to fit the tolerance level of all drug tolerant mutants with a single unified proteomic model. This included several cycles of feature reduction to reach a small number of marker proteins indicative of tolerance physiology (Figure 2a). Whole proteome analyses of drug tolerant isolates identified 42’104 peptides that were assigned to 3’487 proteins and 11’021 annotation terms (*e.g.*, GO/COG terms, domain and structure annotations). To identify proteome features from this large dataset that covary with tolerance, we employed a simple machine learning approach called projection on latent structures regression (plsr) (Figure S2a) ^34^. plsr identifies complementary hyperplanes that best sketch data along covarying parameters (*e.g.*, tolerance). These multivariate axes rely on specific data subsets that can then be exploited to build predictive models. To be able to better interpret the biological patterns exposed by the models while maintaining prediction accuracy, we streamlined model complexity by coupling plsr analyses to iterative cycles of feature reduction. While initial models were computed from the entire dataset, only the most predictive variables were considered for computation of the next cycle to gradually assemble simplified models with the most relevant proteomic variables (Figure S2b,c). The predictive power of the models was tested through leave-one-out cross-validation of the level of survival upon treatment with tobramycin or ciprofloxacin. The most accurate models for both antibiotics exploited similar parts of the initial dataset, sharing roughly half of the biomarkers identified, indicating that common physiological changes drive tolerance against different classes of antibiotics (Figure 2b).

**Figure 2:**
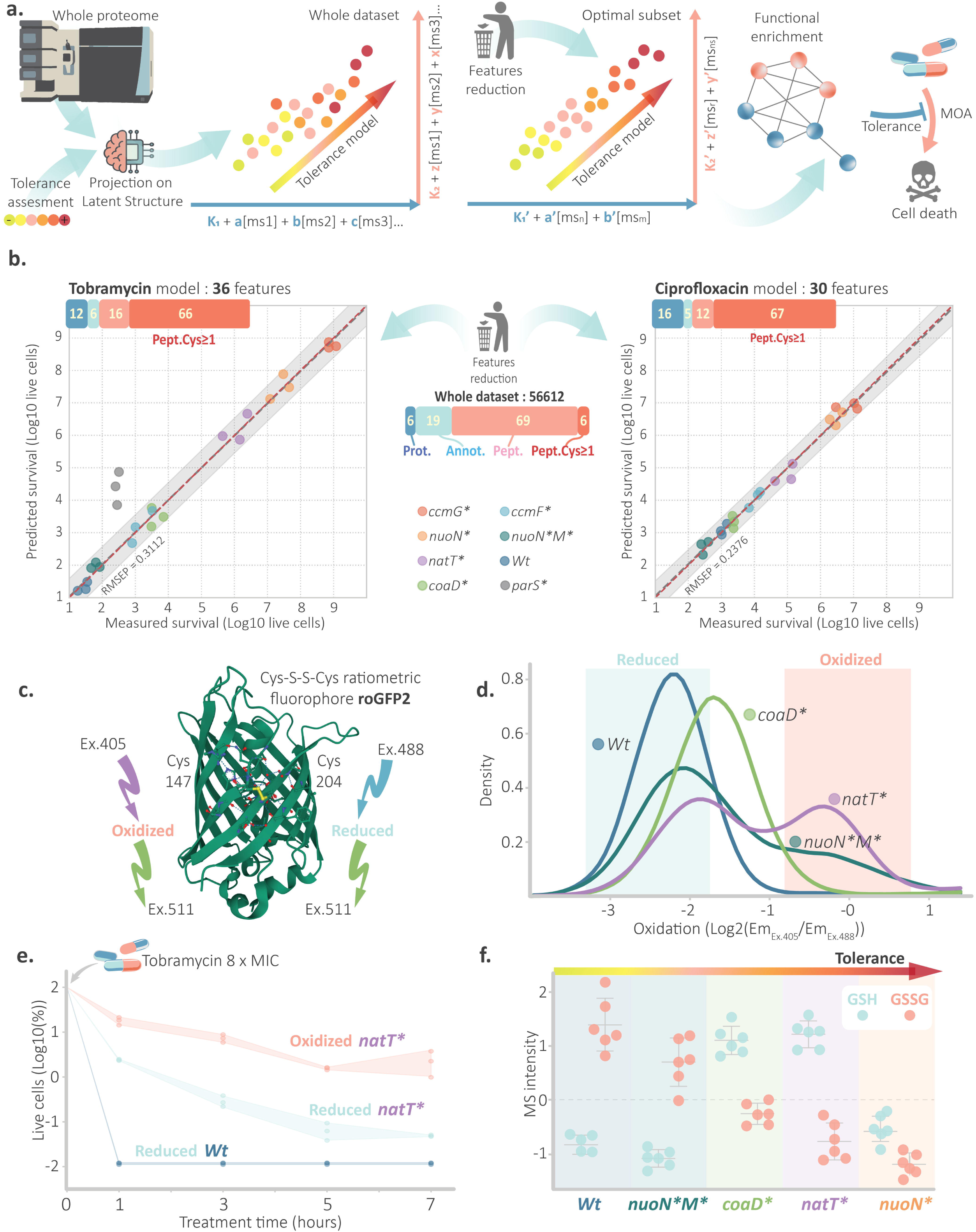
Cysteine oxidation is a biomarker for antimicrobial tolerance. **(a)** Overview of the workflow implementing a projection to latent structures regression (plsr) to predict antimicrobial tolerance based on proteomic data. The model is iteratively refined by feature reduction to identify an optimal subset of predictors. Feature reduction allows for functional enrichment analyses and elucidation of potential mechanisms of action (MOA), linking protein features with tolerance phenotypes. **(b)** Leave-one-out cross-validation performance of plsr models for predicting survival following exposure to tobramycin (left) or ciprofloxacin (right). Predicted and experimentally observed survival of individual strains is shown with an inoculum of 10^9^ cells, and 3 h of treatment with tobramycin (16 µg/mL) or ciprofloxacin (2.5 µg/mL). Linear regression (red dotted lines), x=y function (grey dotted lines) and 95% confidence intervals (shaded areas) are indicated. Bar plots summarize molecular categories of features retained in the final models and for the overall dataset (center): peptide (red/pink), protein (dark blue) or annotation (light blue) features. Peptide-level features dominated the predictive sets with Cys-containing peptides (red) significantly enriched among predictors datasets. **(c)** Schematic of the redox-sensitive biosensor roGFP2, which reports cysteine oxidation via ratiometric fluorescence (www.rcsb.org/3d-view/1JC1/1). The sensor discriminates between reduced (Cys thiols) and oxidized (disulfide bond) states through differential excitation at 405 nm and 488 nm, with emission at 511 nm. **(d)** Flow cytometry-based quantification of cysteine oxidation using roGFP2 expressed in low-tolerance (WT, blue; *nuoN*M\**, turquoise) and high-tolerance strains (*natT\**, purple; *coaD\**, green). Density distributions (Y-axis, LOESS regression, n=3) reflect single-cell heterogeneity of redox status with reduced (blue) and oxidized (red) subpopulations indicated. **(e)** Survival of subpopulations of the *P. aeruginosa natT\** hyper-tolerant strain sorted from panel (d) during treatment with tobramycin (16 µg/mL). Single replicates (n=3) (dots), LOESS regressions (lines) and 95% confidence intervals (colored areas) are shown. **(f)** Reduced (GSH, blue dots) and oxidized (GSSG, pink dots) glutathione in strains with increasing tolerance as determined by mass spectrometry. Averages and standard deviations from mean are indicated (n=6).

The best predictions for tobramycin tolerance required only 36 features, including 26 peptides, four proteins and eight domain annotations (Figures 2b, S2d,e). Annotations and proteins features exploited by the model corresponded to the ribosomal protein L36 and to three homologs of the universal stress protein A (PA4328, PA1789 and PA5027), which is involved in the response to oxygen limitations ^35^. The 26 peptides matched to 25 individual proteins, 13 of which were classified as “biosynthetic processes” (GO:0044249, false discovery rate: 0.000134). Strikingly, more than 80% of these peptides contained at least one cysteine, compared to only 7.7% of Cys-containing peptides in the entire data set (Figure 2b, S2d,e). Similarly, the best model for ciprofloxacin tolerance relied on 30 proteomic markers, including primarily cysteine-containing peptides and the universal stress protein UspA (PA4328) (Figures 2b, S2d,e). The only hyper-tolerant strain, which the proteome-based model was not able to accurately predict was *parS\** (Figure 2b, left). Mutations in the ParS sensory protein are known to generally enhance antibiotic survival by limiting drug uptake and stimulating drug efflux ^31,36^. Accordingly, the MexXY efflux system was significantly upregulated in the *parS*\* strain (Figure S3), indicating that its tolerance mechanism differs from the other hyper-tolerant strains analyzed.

### Cysteine oxidation accurately predicts antibiotic tolerance

Because Cys-containing peptides—but not other peptides from the same proteins—were selectively enriched in hyper-tolerant strains, we hypothesized that cysteine modifications influence mass spectrometry detection in ways predictive of tolerance. Thiol groups of cysteines are highly reactive and can reversibly oxidize to form disulfide bridges with other thiols or form stable intermediates such as sulfinic, sulfenic or sulfonic acids ^37^. To monitor the redox state of cysteines in individual bacteria, we used roGFP2, a redox-sensitive fluorescent reporter that relies on the dynamic oxidation of two surface-exposed Cys residues (Figure 2c) ^38,39^. Because some hyper-tolerant mutants were prone to genetic suppression during genetic manipulation, our analyses focused on *natT\**, *coaD\** and *nuoN*-nuoM* mutants. Strikingly, tolerant strains displayed broadened roGFP2 spectra in stationary phase, consistent with increased cytoplasmic oxidation under tolerance-inducing conditions (Figure 2d). To test if increased oxidation correlates with drug tolerance at the single cell level, stationary cultures of wild type and *natT*\* expressing roGFP2 were sorted into reduced and oxidized fractions and exposed to high concentrations of tobramycin. While all wild-type cells were in a reduced state and were efficiently killed, almost 10% of the oxidized *natT*\* subpopulation survived 3 hours of treatment and ∼3% survived even prolonged exposure to high concentrations of tobramycin (Figure 2e). By contrast, survival of the reduced *natT*\* subpopulation was ∼100-fold lower, indicating that up to 99% of persisters resided in the oxidized fraction.

In line with these observations, metabolomic analyses of drug tolerant stains revealed altered ratios of reduced (GSH) versus oxidized glutathione (GSSG), with increasing tolerance associated with a progressive shift toward oxidative physiologies (Figure 3d). Upon experiencing redox stress, *P. aeruginosa* over-compensates by biasing its primary redox buffer glutathione toward the reduced form ^40^, explaining why GSH/GSSG ratios rose in parallel with tolerance. Accordingly, proteomic open searches for post-translational modifications revealed an accumulation of peptides containing intramolecular disulfide bridges reflecting the extent of oxidative stress in tolerant strains (Figure S4a). In contrast, the observed decrease in GSSG and in peptides with glutathionylated cysteines (which protect thiols from irreversible over-oxidation to sulfonic acids ^41^) likely reflects a stimulated anti-oxidative response in these strains (Figure S4b). Together, these findings identify endogenous thiol oxidation and the accompanying homeostatic adjustments as robust indicators of antimicrobial tolerance.

**Figure 3:**
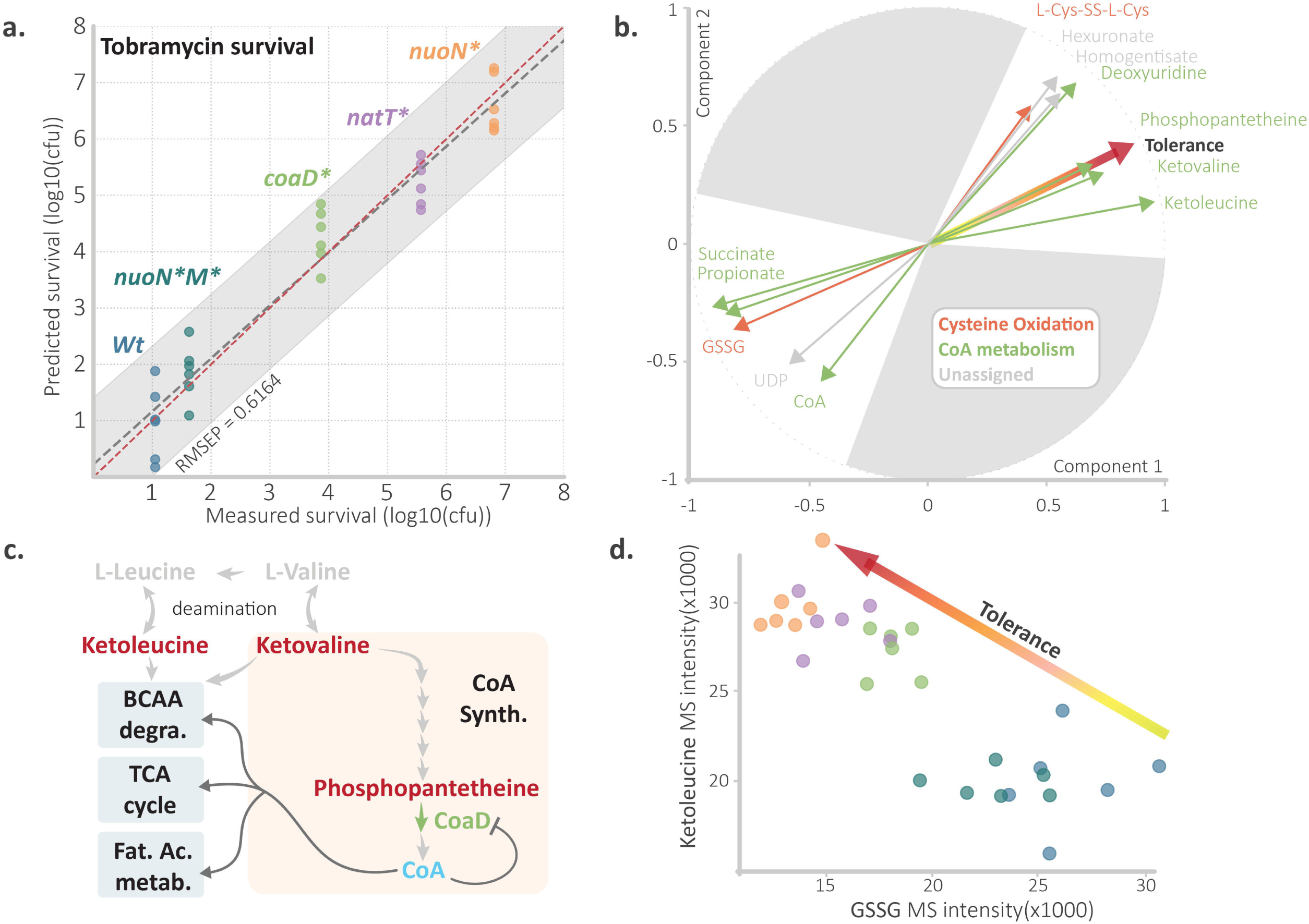
Thiol oxidation and Coenzyme A depletion predict drug tolerance. **(a)** Prediction of survival of *P. aeruginosa* strains based on metabolomics-derived plsr model (leave-one-out cross-validation). Predicted (Y-axis) and experimentally determined (X-axis) survival during treatment with 16 µg/mL tobramycin are shown as loglJlJ CFUs. Each point represents one experimental replicate derived from an initial inoculum of 10lJ cells. Linear regression (red line), x=y function (grey line) and 95% confidence intervals (shaded area) are shown. **(b)** Contribution of metabolites for the prediction of antibiotic tolerance in a biplot representation of the full metabolomic plsr model. The yellow-to-red gradient arrow denotes the tolerance vector, indicating direction and magnitude of correlation with antibiotic tolerance. Green (CoA related), orange (thiol oxidation related) or grey (unassigned pathways) arrows indicate metabolite markers that are strongly correlate or anticorrelate with tolerance. Collinearity with the tolerance vector indicates high predictive relevance. Grey-shaded regions correspond to markers with negligible or non-aligned contribution to tolerance. **(c)** Schematic of Coenzyme A (CoA) biosynthesis with enzymatic steps indicated by grey arrows or in green for phosphopantetheine adenylyl transferase (CoaD). Components that are strongly accumulated (red) or depleted (blue) in hyper-tolerant strains are highlighted. Feedback inhibition of CoaD by CoA is indicated. Major metabolic roles of CoA in TCA cycle, fatty acid metabolism and branched chain amino acid catabolism (BCAA degra.) are indicated. **(d)** A combination of a CoA biosynthesis (ketoleucine) and a thiol oxidation (oxidized glutathione, GSSG) marker is sufficient to discriminate susceptible and drug tolerant strains. The scatter plot shows GSSG (X-axis) and ketoleucine (Y-axis) levels as determined by mass spectrometry in untreated populations of different hyper-tolerant strains of *P. aeruginosa*. Individual replicates (dots) are colored as in panel (a).

### Antibiotic tolerance correlates with coenzyme A limitations

To investigate how thiol oxidation relates to antibiotic tolerance, we next established metabolomic profiles of strains with varying levels of tolerance (Table S1). Models were built from 252 metabolites and were optimized through feature reduction as outlined above. Only 13 metabolites were required to predict tolerance with root mean square error of prediction (RMSEP) as low as 0.616 (Figures 3a; S5). Strong predictors for tolerance included phosphopantetheine, ketovaline, ketoleucine, deoxyuridine, homogentisate, hexuronate or free Cys disulfides all of which correlated positively with the tolerance. In contrast, metabolites such as oxidized glutathione (GSSG), succinate, propionate, UDP and CoA showed strong anti-correlations (Figure 3b). Tolerance also correlated with other oxidation indicators such as deoxyuridine ^42^ or the pyomelanin component homogentisate ^43^ (Figure 3b). Intriguingly, several intermediates of CoA biosynthesis (ketovaline, ketoleucine and phosphopantetheine) accumulated in drug tolerant strains, while CoA itself was strongly reduced (Figure 3b,c). A single metabolic marker (ketoleucine accumulation) in combination with a marker for thiol-oxidative stress (GSSG) was sufficient to predict survival rates (Figure 3d). CoA depletion together with the observed accumulation of CoA precursors identified the phosphopantetheine adenylyl transferase CoaD as a potential bottleneck in CoA biogenesis of drug tolerant strains ^30^ (Figure 3c). Because these metabolic changes did not correlate with different CoaD levels (<u>massive.ucsd.edu:</u> MSV000091240, Figure S3), we hypothesized that CoaD activity is compromised in hyper-tolerant strains. CoaD catalyzes the essential penultimate step in CoA biosynthesis and also contributes to pathway control through feedback inhibition by the end-product CoA ^30^. Notably, one of the hyper-tolerant mutants analyzed harbors a mutation in *coaD* (*coaD\**), further implicating this enzyme as a key node in antibiotic tolerance. Given its pivotal role in central metabolic pathways, including the tricarboxylic acid (TCA) cycle, fatty acid β-oxidation, and branched-chain amino acid (BCAA) catabolism, CoA limitations could account for the observed reduction in succinate and propionate levels, as well as the accumulation of deaminated BCAA degradation intermediates such as ketovaline and ketoleucine in tolerant strains ^44–47^.

To corroborate the link between CoA limitations and antibiotic tolerance, we examined tolerance and proteome composition of the *nuoN\** strain across different growth stages. Survival of *nuoN\** during antibiotic exposure increased progressively over five orders of magnitude as cultures approached stationary phase (Figure 4a). To explore the underlying physiology, we generated comprehensive proteomic data sets of *nuoN\** across growth phases and used them to construct plsr models. These models accurately predicted survival (RMSEP = 0.3028) based on 200 proteomic variables representing 65 proteins (Figure 4b). A STRING-based STITCH platform analysis, which integrates protein and metabolic data ^48^, revealed strong interconnections among these proteins (PPI enrichment *p-value*: 2.76e-09), suggesting that they collectively define a growth phase–dependent tolerance pathway (Figure 4c). Strikingly, a large fraction of the proteins were directly (43%) or indirectly (18%) linked to CoA in *P. aeruginosa* and other species. Associated pathways included branched chain amino acid (leucine, isoleucine and valine) and ketone body degradation, TCA cycle and general carboxylic acid metabolism. The signature also comprised proteins involved in translation, and the universal stress protein UspA. This is in line with a general involvement of stress proteins during CoA depletion and with the view that translation arrest contributes to treatment survival ^15^.

**Figure 4:**
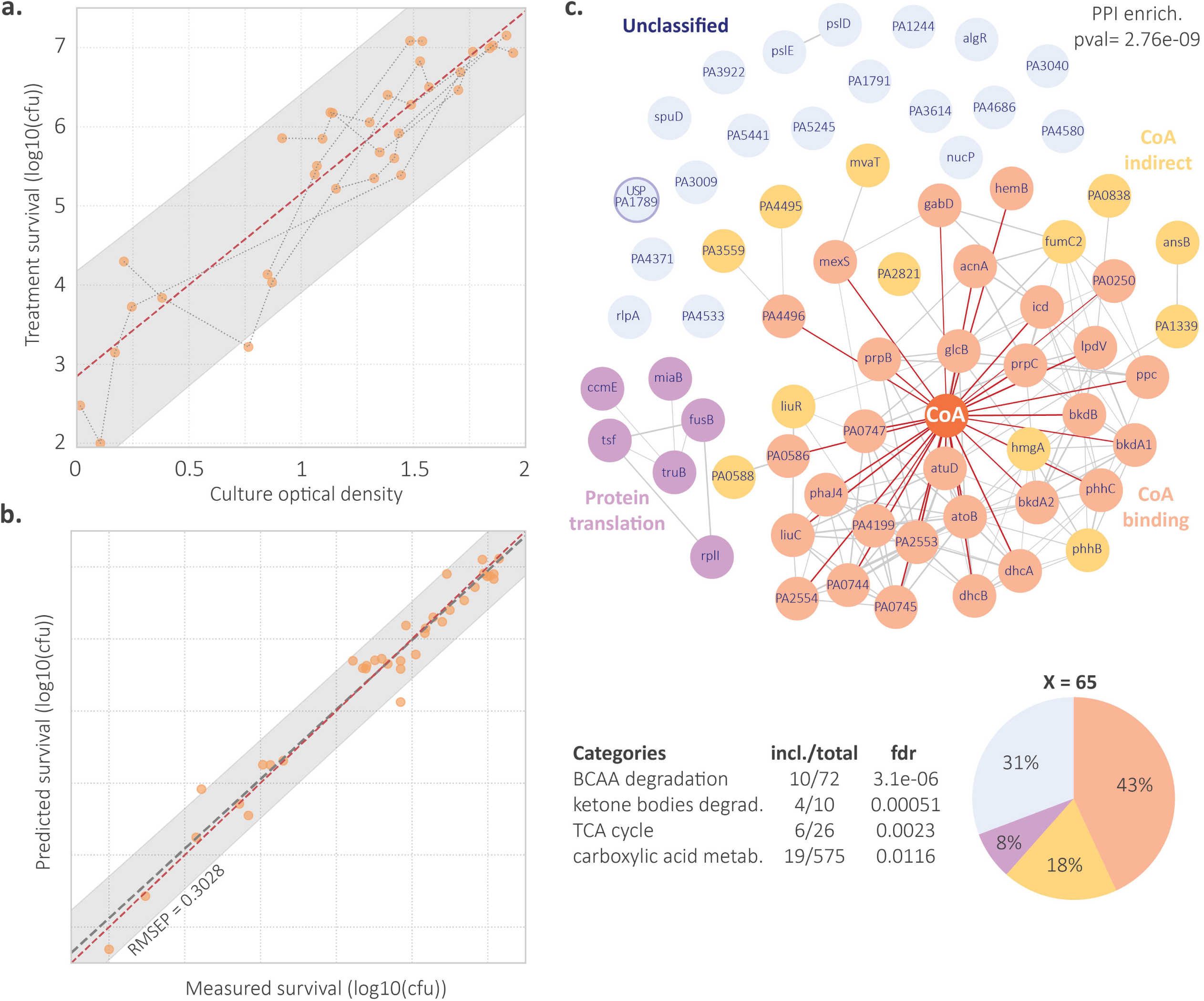
A proteomic model predicts growth-phase dependent increase of tolerance. **(a)** Survival of the *nuoN\** hyper-tolerant strain correlates with growth phase. Experimentally determined survival (loglJlJ CFU; Y-axis) after exposure to tobramycin (3 h; 16 μg/ml) is plotted against the optical density (ODlJlJlJ, X-axis) of a growing culture of *nuoN\** at the time of treatment. Individual measurements of a given culture are connected by grey dotted lines. Linear regression (red dotted line) and the 95% confidence interval (grey area) are indicated. **(b)** Leave-one-out cross-validation of a proteomics-based plsr model predicting tobramycin survival of the *nuoN*\* hyper-tolerant strain harvested at different phases of growth as indicated in (a). The predicted survival (Y-axis; loglJlJ CFU) is plotted against experimentally determined survival (X-axis; loglJlJ CFU) as shown in panel (a). Linear regression (red dotted line), x=y function (grey dotted line) and 95% confidence interval (grey area) are shown. The optimal uses 200 variables corresponding to a subset of 65 proteins. **(c)** Functional interaction network of 65 proteins identified as predictive for survival of *nuoN\** at different growth stages as shown in (a) and (b) using the STRING-based STITCH platform (http://stitch.embl.de). Protein–protein interaction (PPI) enrichment analysis revealed significant connectivity (blue edges; p = 2.76 × 10lJlJ) with dominant clusters related to CoA (CoA binders: salmon; indirect interactors: orange) and protein translation (purple). Unclassified proteins (steel blue) include UspA (highlighted by blue circle). The distribution of functional categories of predictive proteins is shown as pie chart with pathway enrichment parameters and false discovery rates [fdr]) listed for components of branched-chain amino acid (BCAA) degradation, ketone body degradation, TCA cycle, and carboxylic acid metabolism.

Together, these experiments identify shortages of coenzyme A as a potential indicator for the ability of *P. aeruginosa* to survive antibiotic exposure.

### Coenzyme A shortages mediate drug tolerance

CoA carries a reactive thiol group that normally serves to generate activated CoA thioesters. Although relatively resistant to oxidation, CoA can form homo- or mixed disulfides under oxidative conditions, potentially reducing the pool of free, active CoA ^49^. Mixed CoA disulfides (*e.g.* CoA-S-S-Cys-protein or CoA-S-S-glutathione) were shown to accumulate in bacteria ^50,51^ and were proposed to silence metabolism during *Bacillus* sporulation ^52,53^ or reversibly inhibit key enzymes such as RNA polymerase ^54^ and glyceraldehyde-3-phosphate dehydrogenase ^51–53^. To test the causal relationship between CoA activity and antibiotic tolerance, we first demonstrated that CRISPR-cas-mediated knock down of *coaD* increased survival of *P. aeruginosa* (Figure 5a). Conversely, over-expression of *coaD* reduced survival of the hyper-tolerant *natT\** strain by more than 20-fold (Figure 5a). Since *P. aeruginosa* CoaD is subject to complex feedback regulation ^30^, we replaced it with the orthologue from *Staphylococcus aureus,* which we hypothesized would function robustly under fluctuating CoA pools. Unlike *P. aeruginosa* and other gram-negative bacteria, *S. aureus* does not rely on glutathione for redox homeostasis, but instead maintains high intracellular CoA levels to buffer redox changes ^55,56^. Expression of *S. aureus coaD* reduced the survival of the *natT\** strain by ∼100-fold (Figure 5a) and led to an increase of cellular concentrations of CoA, biotin, phosphoglycolate, succinate and cytidine (Figure 5b; Table S2), indicating that restored CoA levels boosted CoA-dependent biotin biogenesis, succinate production and the TCA cycle ^57^. Because *S. aureus* critically depends on its reduced CoA pool for redox balance, it encodes a CoA disulfide reductase (CoADR_st_) that restores functional CoA by reducing oxidized CoA thiols ^49,58^. Intriguingly, expression of *coADR_st_* also reduced survival of the *natT\** strain by ∼100-fold (Figure 5a). Although CoA levels remained unchanged under these conditions, elevated cytidine and biotin suggested that CoA-dependent pathways were at least partially restored (Figure 5b). Moreover, in line with CoADR_st_ utilizing FADH_2_ as reducing power for CoA disulfide reduction ^58^, we observed robust accumulation of oxidized flavin adenine dinucleotide (FADox) under these conditions.

**Figure 5:**
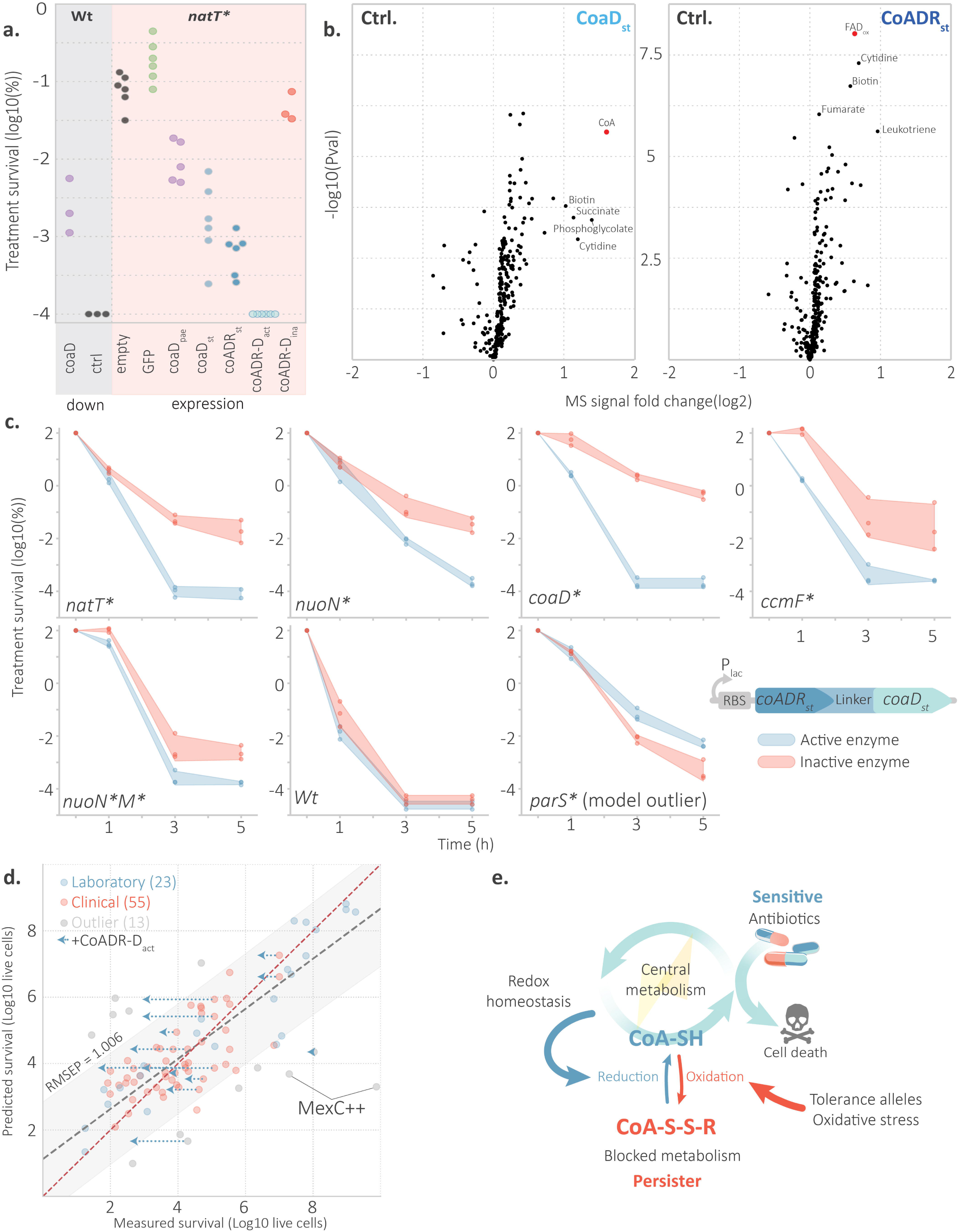
Coenzyme A limitations as major driver of *P. aeruginosa* tolerance. **(a)** Survival of *P. aeruginosa* upon modulation of CoA levels. Survival of *P. aeruginosa* wild type (grey area) upon dCas9-mediated knockdown of *P. aeruginosa coaD* with (purple) and without (grey) expression of a *coaD*-specific sgRNA. Survival of the hyper-tolerant strain *natT\** upon expression of different *coaD* and *coADR* variants (pink area). The *natT*\* contained an empty plasmid control or plasmids expressing *gfp* (green), *coaD_pae_*from *P. aeruginosa* (purple), *coaD_st_* from *S. aureus* (steel blue), *coADR_st_* from *S. aureus* (blue) or an engineered fusion of *coaD_st_* and *coADR_st_* (*coADR-D_st_*) wild-type sequences (_act_; light blue) or inactive mutant versions (_ina_; red). The fraction of cells (Log_10_) surviving three hours of treatment with tobramycin (16 μg/ml) is shown (n=3-6). **(b)** Metabolomic analyses of *natT\** expressing either *coaD_st_* or *coADR_st_*in stationary phase (when tolerance is scored). Volcano plots depict 259 metabolites enriched (Log2FC > 0) or depleted (Log2FC < 0) compared to the same strains with a mock plasmid expressing *gfp*. The Y-axis indicates reproducibility (-log10 of the p-value). Red dots indicate the direct products of enzymes expressed. **(c)** Expression of the CoADR-D fusion protein (lower right) resensitizes hyper-tolerant *P. aeruginosa* strains. Tobramycin-mediated killing (16 μg/ml) is shown for strains expressing an active (blue) or an inactive (red) form of CoADR-D. Fraction of surviving cells (Y-axis, Log_10_) are shown as individual dots (n=3) with standard deviations shown as colored areas. Expression of the *coADR*-*D* genes is controlled by a lactose inducible promotor (P_lac_) with a synthetic RBS maximizing translation. **(d)** Prediction of survival rates of clinical isolates based on Cys-peptides. Survival of 68 drug sensitive (MIC tobramycin ≤ 4 & MIC ciprofloxacin ≤ 1) clinical isolates (red dots) was determined experimentally (tobramycin 16 μg/ml; 3 h) (n=3) (X-axis) and predicted based on proteome analyses of the same strains prior treatment (massive.ucsd.edu: MSV000091240) (Y-axis). The model was trained with proteome data from clinical isolates and lab strains with different levels of tolerance (blue). Linear regression (red dotted line), x=y function (grey dotted line) and 95% confidence interval (grey area) are shown. Re-sensitization of drug-tolerant clinical strains expressing *coADR-D*_act_ is indicated by blue dotted arrows. Some model outliers with high levels of MexC efflux pump are indicated. **(e)** Model of CoA-mediated antimicrobial tolerance. Physiological levels of reduced CoA maintain an active central metabolism, ensure redox homeostasis and promote antibiotic susceptibility (blue arrows). We propose that moderate oxidative stress enhances the oxidation of thiols including CoA (red arrows), which in turn limits the availability of free CoA. This temporarily blocks central metabolic pathways, thereby leading to drug tolerance.

Since CoaD_st_ contributes to the overall CoA pool whereas CoADR_st_ maintains its bioactivity, we next asked whether their functions would display additive effects in restoring drug susceptibility. To test this, we engineered a fusion protein consisting of full-length CoaD_st_ and the nucleotide-disulfide oxidoreductase domain of CoADR_st_ (Figure 5c) ^58^. Expression of the CoaD-CoADR fusion completely abolished tolerance of the *natT\** strain (Figure 5a) and enhanced killing of most other drug-tolerant strains in a MIC-independent manner (Figure 5c; Figure S6). Introducing active site mutations in the CoaD and CoADR moieties of the CoaD-CoADR fusion abolished drug resensitization (Figure 5a,c), demonstrating that the observed effect requires catalytic activity. Notably, expression of the CoaD-CoADR fusion did not reduce drug tolerance of the efflux pump over-expressing *parS\** mutant, verifying that the activity of the fusion protein specifically counteracted CoA-based tolerance mechanisms.

### A redox-based model predicts antibiotic tolerance in clinical *P. aeruginosa* isolates

Although drug-tolerant isolates of *P. aeruginosa* exhibit similar tolerance *in vitro* and *in vivo* ^59^, tolerance mechanisms that evolve *in vitro* may differ fundamentally from those arising during human infections ^2,4,60^. To address this, we asked whether the redox-based tolerance model is able to predict tolerance levels of clinical *P. aeruginosa* isolates. For this, we analyzed the proteome of 68 strains recovered from chronically infected airways, which had previously been shown to display increased tolerance ^2^. Unlike the collection of hyper-tolerant *P. aeruginosa* strains evolved *in vitro*, clinical isolates display substantial genetic heterogeneity. This complicates proteome analysis because genetic variations and differences in protein expression can impair peptide detection. To mitigate genetic heterogeneity, we trained a new reductive plsr model based on ratios between cysteine-containing peptides (oxidative marker) and cysteine-free peptides from the same proteins (internal expression standards). This allowed cysteine oxidation levels to be assessed independently of inter-strain variation. The resulting optimized model incorporated 18 peptide ratios from 11 proteins across proteomes of 55 clinical and 7 laboratory strains. The model accurately predicted tolerance levels of 80% of the isolates, with accuracies comparable to those obtained for tolerant strains evolved *in vitro* (Figure 5d). The 11 proteins selected by the model (Table S3) are conserved and function in core metabolic processes (enrichment FDR:0.003) including DNA repair, TCA cycle, fatty acid metabolism, amino acid and nucleotide biosynthesis, transcription, translation, glycolysis/gluconeogenesis and oxidative phosphorylation. The unbiased selection of markers spanning distinct pathways suggests that tolerance involves the concomitant perturbation of multiple essential cellular processes.

Thus, thiol-based models that are able to predict drug tolerance evolved *in vitro* also performed well in predicting tolerance of clinical isolates, supporting the view that increased thiol oxidation is a fundamental mechanism underlying antimicrobial tolerance during chronic infections. This conclusion is further supported by the finding that expressing the *S. aureus* CoaD-CoADR fusion protein significantly reduced survival of a random selection of drug tolerant clinical isolates during tobramycin treatment (Figure 5d). Of the 68 patient isolates analyzed, only 13 were not accurately predicted and were excluded as outliers from the final dataset. Notably, several outliers underestimated by the model displayed elevated expression of the inducible efflux pump component MexC ^61^ (Figure 5d, massive.ucsd.edu: MSV000091240). Efflux pump upregulation is known to confer broad antimicrobial tolerance ^62^, providing a plausible explanation for why tolerance in these isolates was poorly captured by the model.

## DISCUSSION

Our studies show that hyper-tolerant strains of *P. aeruginosa* exhibit shared physiological traits that promote oxidation during nutrient depletion. A consistent theme emerging is that mild thiol oxidation underlies this primed tolerant state. While reactive oxygen species (ROS) are classically associated with antibiotic lethality—through unbalanced respiration, NADH accumulation, and oxidative bursts during treatment ^26,63–65^—our results reveal that more subtle oxidative changes can have the opposite effect, favoring survival. Specifically, thiol oxidation correlated consistently with tolerance across strains, whereas responses mediated by other oxidative stress regulators (*e.g.*, *soxR*, *oxyR*), greatly varied across different isolates (Figure S3; MassIVE : MSV000091240) ^66,67^.

Mechanistically, this protective oxidative state converges on the thiol-molecule CoA. Tolerant strains displayed markedly reduced levels of CoA, while restoring active CoA abolished tolerance (Figure 5). Since the level of several intermediates upstream of the CoaD-catalyzed step of CoA synthesis correlated with tolerance, CoaD emerges as a central bottleneck. Oxidized CoA disulfides, while catalytically inactive, may still exert feedback inhibition on CoaD as described for free CoA ^30,68,69^, thereby depleting active CoA while simultaneously blocking its replenishment. Consistent with this model, heterologous expression of a feedback-insensitive CoaD from *S. aureus* increased CoA levels and strongly reduced tolerance (Figure 5). Alternatively, CoaD inhibition could arise from direct thiol modifications such as protein coalation, which were recently linked to oxidative stress adaptation ^70^. Notably, our unbiased approach did not identify other growth rate–limiting proteins (*e.g.* ribosomes) or metabolites (*e.g.* ATP), as indicators of tolerance. This is consistent with the notion that, despite the overall correlation between slowed growth and tolerance, dormancy alone is not sufficient to explain bacterial drug survival ^71,72^.

Oxidative stress emerges as a double-edged sword: at high levels, respiratory hyper-oxidation amplifies antibiotic killing, whereas at milder levels, it primes cells for dormancy and persistence ^73–76^. In line with this, microscopy tracking of individual lineages upon drug treatment highlighted that, paradoxically, susceptible cells killed more rapidly during treatment are more difficult to discriminate from persister lineages than cells dying at a later time point (Figure S1c). Their physiological traits diverged from persisters only for those features that were recorded close to drug treatment (Figure S1d,e). This implies that lineages with a shared history in stationary phase diverge into opposite fates, persistence or drug hyper-sensitivity and rapid death. Thus, CoA-oxidation may not enforce a uniform outcome, but instead generate heterogeneous cell fates that either promote survival or accelerate killing. We postulate that CoA oxidation functions as a reversible “metabolic fuse” that keeps the respiratory chain from collapsing under oxidative pressure. By sequestering CoA into disulfide forms, catabolic activities and electron flow would be transiently curtailed, thereby reducing respiratory pressure. Once the redox balance is restored, the relatively low potential energy of thiol disulfides allows their reversible reduction, enabling growth resumption after a period of metabolic dormancy—similar to CoA-dependent spore germination in *Bacillus* ^52,53^.

CoA-based tolerance mechanisms may be wide-spread and of general nature. Indeed, similar physiologies have been reported in unrelated organisms. Small colony variants recurrent in chronic and relapsed *S. aureus* infections often carry electron transport chain mutations that confer tolerance to diverse antibiotics ^77,78^ and up-regulation of ROS detoxification genes ^79^. Recently, *Salmonella enterica* persisters in macrophages were shown to rely on oxidative stress triggered by host-derived reactive nitrogen species, which ultimately cause an arrest of the central metabolism ^80^. Similarly, proliferating cancer cells elevate ROS through alternative metabolic programs ^81^, while simultaneously boosting antioxidant defenses to withstand chemotherapy ^82–84^.

Altogether, this work provides mechanistic insight into antibiotic tolerance in an important human pathogen. By causally linking stationary-phase physiology, respiratory chain function, and CoA-centered redox control, we outline a generalizable framework for persistence. These findings not only identify biomarkers and diagnostic tools for tolerance but also highlight molecular targets—such as CoaD—for therapeutic intervention against antibiotic tolerance during chemotherapy.

## Material and methods

### Bacterial strains and culture conditions

Strains used in this study are listed in Table S4 in the supplemental material. Unless otherwise stated, *P. aeruginosa* PAO1 and all *E. coli* strains were grown at 37°C in Luria-Bertani (LB) medium^85^ with shaking at 170 rpm or, alternatively, statically and solidified with 1.3% agar when appropriate. Knock-down and plasmid expression based assays (Figure 5abc, growth and killing kinetics) were performed in minimum medium MOPS medium ^86,87^ supplemented with 20 mM succinate. Tetracycline was used for plasmid maintenance at 100 μg/ml for *P. aeruginosa* and at 12.5 μg/ml for *E. coli*.

### Plasmids and oligonucleotides

Plasmid used in this study are listed in Tables S4 and synthetized genes and primers in Table S5.

### Molecular biology procedures

Cloning was carried out in accordance with standard molecular biology techniques. pME6032-*px2::grx1::roGFP2* has been engineered from a pME6032 derivative were the lac promotor has been replaced by the constitutive px2 promotor ^88^ ligated with the PCR products from pSRK-*grx1*::*roGFP2* ^89^ (oligos UJ.14013 and UJ.14014). pME6032–*coaD_Pae_*, was engineered from ligations between pME6032 (Table S4) and the PCR product of the coaD gene from PAO1 (PA0363) (oligos UJ.16234 and UJ.16235). pME6032–*coaD_st_*, pME6032–*coADR_st_*, pME6032–*coADR-D_act_*, and pME6032–*coADR-D_ina_* and pME6032–*gfp* were engineered from Gibson ligations ^90^ between pME6032 restricted with Eco53KI (Table S4) and the PAO1 codon-optimized synthetic DNA (Twist Bioscience, San Francisco, US) encoding the different genetic targets (Table S5). Maximized RBS ^91^ and pME6032 overhanging regions around the unique site Eco53kI are included in these sequences. Knock down strains are derivatives from a Dcas9 expressing variant of PAO1 *natT*\*: the *dCas9* gene was integrated in the chromosome as described elsewhere ^92^ using the pUC18-mini*-Tn7T-Lac-dCas9* followed by removal of the gentamicin marker using the pFLP2 as previously described^93^. The plasmid carrying the sgRNA, pBx-Spas-sgRNA, was modified by removal of all additional BasI sites and exchange of the BbsI for BsaI sites using QuickChange and GoldenGate cloning, resulting in pBx-Spas-sgRNA2. GoldenGate cloning was then used for integration of the desired sgRNA into pBx-Spas-sgRNA2.

### Antibiotic survival assays

To measure survival under antibiotic treatments, overnight cultures, each grown from a single colony in liquid medium, were diluted to an OD of 0.1 into fresh medium supplemented with a tobramycin concentration of 16 µg.ml^-1^ or ciprofloxacin 2.5 µg.ml^-1^. At the time points indicated in the figures, aliquots of the cultures were sampled, diluted, and plated on LB medium plates. CFU were counted after overnight incubation and over a 2-day period to make sure that potentially delayed CFU had appeared.

### Proteome analyses

Mass spectrometry samples were collected from cultures used for the antibiotic survival assays (right before the start) in order to maximize the efficiency of the regression approach. Thus replica variations may also contribute to fine tuning of models. Sample generation is detailed elsewhere ^34^. In brief, the equivalent of 1 mL at OD600 0.6 (corresponding to ca. 5 × 10^8^ CFU/mL) was collected, spun down (10 000 rcf, 2 minutes), supernatant was removed, pellets were flash frozen in liquid nitrogen and stored at −80 °C. For cell lysis prior protein extraction, samples were sonicated and heated in MS compatible lysis buffer. Chloroacetamide was added to the samples to allow proper cysteine identification. Sample pH was adjusted between 8 and 9 before starting an overnight trypsin digestion at 37 °C overnight without shaking. Peptides were then solid-phase purified using SDB-RPS extraction cartridges. After elution, peptide mixtures were concentrated under vacuum to complete dryness and stores at −80 °C. For LC-MS analysis, peptide concentration was adjusted to 0.5μg/μL using LC buffer A. For each sample, 1μg of total peptides was subjected to LC-MS analysis using a dual pressure LTQ-Orbitrap Elite mass spectrometer connected to an electrospray ion source and a column heater set to 60 °C. Peptide separation was carried out using an EASY nLC-1000 system equipped with a RP-HPLC column (75μm × 30 cm) packed with C18 resin. The HPLC was run as successive linear gradients from 95% solvent A and 5% solvent B to 35% solvent B over 50 min, to 50% solvent B over 10 min, to 95% solvent B over 2 min and stay at 95% solvent B over 18 min at a flow rate of 0.2μL/min. The data acquisition mode was set to obtain one high resolution MS scan in the cyclotron part of the mass spectrometer: Resolution at 120,000 full width at half maximum (400 m/z, MS1) followed by MS/MS (MS2) scans in the linear ion trap of the 20 most intense MS signals. The charged state screening modus was enabled to exclude unassigned and single-charged ions and the dynamic exclusion duration was set to 30 s, the collision energy to 35%, and the number of microscans acquired for each spectrum to one. “.raw” files were imported into the Progenesis QI software (v2.0, Nonlinear Dynamics Limited) and peptide precursor ion intensities extracted across all samples applying the default parameters. The resulting “.mgf” files were searched using MASCOT against a decoy database with the following criteria: Full tryptic specificity (cleavage after lysine or arginine residues, unless followed by proline), three missed cleavages allowed, carbamidomethylation (on cysteine) set as fixed modification, oxidation (of methionine) and protein N-terminal acetylation accepted as variable modifications, and mass tolerance of 10 ppm (precursor) and 0.6 Da (fragments). Normalized label-free quantification was performed using the SafeQuant R package v.2.3.4 (https://github.com/eahrne/SafeQuant/)^94^ in order to obtain peptides relative abundances. Only isoform-specific peptide ion signals were considered for quantification. Post-translational modifications were exhaustively searched using Msfragger (mass shift open search with default settings) ^95^. Calculations were performed at sciCORE (http://scicore.unibas.ch/) scientific computing core facility at University of Basel.

### Projection on latent structures regression (plsr) and iterative feature reduction

Plsr is a variable regression method based on maximizing the covariance between explicative variables (here, the proteomic data) and the response phenotype (here, survival after drug treatments) (Figure S1abc). A detailed step-by-step procedure is described elsewhere ^34^. Plsr combines different linear regressions over a given data set as depicted in Figure S1a. Plsr purposely exploits different parts of the covariance in each component. This means that the information laying in the data used in the first regression is not used by subsequent regressions. In order to avoid over-fitting, we limited the number of regressions (called components) to two per model. Besides, this allows for the complete representation of generated models by simple 2D-graphics. Performances of models were assessed by leave-one-out cross validation: for each sample a predictive model was computed excluding this sample and the phenotype of this sample was then assessed. The average differences between experimental and predicted tolerance phenotypes were then used as a performance score (i.e., Root Mean Square Error of Prediction, RMSEP) (Figure 2b). Perl and R script templates used for data preparation are available (https://github.com/Pablo-Manfredi/PLS-regression-of-MS-data). These scripts compute the peptide MS signals for each protein and combine peptide and protein MS data with the phenotypic data (e.g., tolerance), perform per sample normalization by dividing each MS signal by the total signal of the sample, execute the first plsr on the whole dataset, run feature selection and output graphics presented in this study. To optimize the biological insight into tolerance physiology, iterative cycles of feature reduction were executed. An initial model is computed from the entire dataset and a score of model contribution is calculated for each variable (*i.e.* variable importance in projection, VIP). VIP assesses the level of contribution of a given variable (here, a peptide, a protein, an annotation or a metabolite) to the current predicting model. Variables with a VIP above 1 are arbitrarily considered for the computation of the next cycle model (*i.e.* another plsr on the data subset). After each cycle, new VIP scores are computed from the new models and a variable selection is operated for VIPs above 1 for integration into the next cycle. This process greatly enhanced the model predictions and best cross-validations were achieved with just few features (*c.a.* 30). Different methods to determine the VIP cutoff were tested without significant improvement over an arbitrary fixed cutoff (=1) for the VIP. This process was repeated until reaching a level where too few variables were left to perform the plsr or when all variables of a cycle subset displayed VIP values above 1 (*i.e.* convergence). Most often, the models with the best predictive power were among the simplest ones (Figure S1b,c and S5). Nevertheless, too minimalistic models proven to be difficult to interpret in terms of biological pathways. Thus, we rather expanded on models that retained a certain number of variables (>30) in order to observe clear functional enrichments. Predictive models based on clinical and laboratory strains were computed in a similar fashion with the difference, for clinical strains, that the input data set was exclusively composed of all the ratios between each cysteine containing peptide detected and each other non-cysteine containing peptides of the same protein.

### Fluorescence activated cell sorting (FACS)

To analyze the thiol-redox physiology of single cells in different drug tolerant mutants, we used the more dynamic variant of the roGFP reporter grx-roGFP2 ^38,96^. For each sample, a single bacterial colony was grown in LB medium for 16 h at 37 °C under agitation (170 RPM). Cultures were sampled for FACS analysis in same circumstances as for antimicrobials killing assays. FACS measurements were performed on a BD FACSAria IIIu Cell sorter. Fluorescences were measured with the following channels Ex488_LP495_BP514/30-H and Ex405_LP502_BP530/30-H. Gates were determined taking into account the non-fluorescent counterparts of analyzed strains (Figure S8).

### Metabolomic analysis

Cultures were sampled for metabolomic analyses in same circumstances as for antimicrobials killing assays. Samples were set at *c.a.* 10^9^ CFU/mL (e.g. 1 mL of OD600 1), pelleted (10 000 rcf, 1 minute) and supernatant was carefully removed. Because of the exceptional sensitivity of mass spectrometers and the versatility of certain metabolites, samples were flash frozen in liquid nitrogen before proceeding with solvent extraction. Each pellet was then resuspended into 0.5 mL of extraction solution (EtOH:H2O, 60:40) by repeatedly pipetting up and down and vortexing. Samples were incubated in a Thermomixer at 75 °C and maximum shaking speed for 1 min before a centrifugation step (10 000 rcf, 30 seconds). The supernatant was collected into a new tube and the extraction procedure was repeated one the remaining pellet. The second extraction was pooled with the first one prior speedvac drying; flash freezing and storage at -80°C. Extracts were analyzed by flow injection – time of flight mass spectrometry on an Agilent 6550 QTOF instrument operated in the negative mode, as previously described ^97^.

### Microfluidic assays

Master molds for the dual-input mother machine, consisting of SU-8 on 4-inch silicon wafers, were constructed by Micro Resist Technology GmbH based on published design files (1 μm cell trap height and 20-μm flow layer height) ^98,99^. Microfluidic devices were then fabricated by pouring Sylgard 184 Polydimethylsiloxane (PDMS, Dow Corning) onto the molds, followed by baking at 80°C for 1 h. Access holes were created using 0.75mm biopsy punchers (Robbins Instruments), and the PDMS devices were bonded to microscope cover slides following plasma treatment and baking at 100°C for 1 min on a hot plate. The devices were connected using PTFE Tubing (0.56mm ID x 1.07mm OD, Adtech) to a pressure pump (Elveflow OB1 MK4). PAO1 LB overnight cultures were centrifuged, filtered twice (25 μm pore-size) and used as spent LB medium (SLB) in the fluidic experiments. To analyze the single-cell activity of the *P. aeruginosa natT\** mutant, experimental setups were divided into different phases: (1) Exponential, (2) Gradual to Stationary, (3) Stationary, (4) Antibiotic treatment, and (5) Recovery. The exponential phase consisted of exposing the cells to fresh LB medium for 3 h. During the gradual introduction to the stationary phase, cells were exposed to increasing ratios of fresh LB and SLB for 2.5 h until the complete exposure to SLB for 10h to mimic stationary phase conditions. During the drug treatment phase, tobramycin diluted in fresh LB medium at a final concentration of 16ug/mL was provided. The experiments ended with a recovery phase of LB medium applied for 24 hrs.

### Microscopy and Image Acquisition

Images were acquired with a Nikon Eclipse Ti2 Inverted Microscope equipped with a Hamamatsu ORCA-Flash4.0 V3 Digital CMOS camera (C13440-20CU) and a 100x NA1.45: Nikon Plan Apo Lambda 100x Oil Ph3 DM objective (MRD31905). Best focusing performance was achieved with the Zeiss low fluorescence Immersion oil for fluorescence-microscopy 518F/37°C, n_e_= 1,518. Florescence imaging was acquired with LED light at 470nm (for GFP) and 575nm (for RFP) using a SPECTRA-X LED Illumination System as light source. Emitted light was filtered using a GFP/mCherry ET dual band Chroma filter (F58-019). Focuse was maintained using the Nikon perfect focus systems and the setup was controlled using the Nikon NIS-Elements software. The sample was maintained at 37°C using an okolab incubator. Images were acquired with intervals of 10 minutes. The data was saved locally in ND2 format.

### Data processing for single-cell segmentation and tracking

Cell segmentation and tracking of single-cells was done using the “Bacteria in Mother Machine Analyzer” (BACMMAN) and its deep-learning algorithm DistNet ^100^. Segmentation and tracking were manually corrected from BACMMAN kymographs using a Wacom tablet. A training and validation datasets were created in BACMMAN and extracted as HDF5 files. Training of DistNet was based on a Colab script provided by BACMMAN (https://colab.research.google.com/gist/jeanollion/d96bc2a8adef58563427932319bdee1b/finetune_d istnet.ipynb). To increase the accuracy of the persister cells segmentation and tracking, data curation and lineage reconstruction was done manually. Lineage reconstruction of susceptible cells was done with a custom-made code (https://github.com/hector-ahg/EMBO_Journal_toxinantitoxin_pseudomonas.git).

### Segmentation and tracking data processing for plsr predictive models

For this analysis, we selected a refined high tracking fidelity dataset comprising 206 persister and 8’423 sensitive lineages with the lowest amplitude of XY variation between consecutive frames. Segmentation and tracking features from frames 33 to 92 (spent medium phase) were extracted and consisted in ’NextDivisionFrame’, ’PreviousDivisionFrame’, ’SizeRatio’, ’GrowthRateSize’, ’SizeAtBirthSize’, ’Size’, ’GrowthRateFeretMax’, ’SizeAtBirthFeretMax’, ’FeretMax’, ’GrowthRateSpinelength’, ’SizeAtBirthSpinelength’, ’Spinelength’, ’GrowthRateSpineWidth’, ’SizeAtBirthSpineWidth’, ’SpineWidth’, ’GrowthRateLocalThickness’, ’SizeAtBirthLocalThickness’, ’LocalThickness’, ’MeanIntensityRFP’, ’MeanIntensityGFP’, ’BoundsXMin’, ’BoundsXMax’, ’BoundsYMin’, ’BoundsYMax’, ’RelativeCoordX’, ’RelativeCoordY’, ’CellPosChannel’, ’DeltaYMin’, ’DeltaYMax’ and the lineage constant values ’num_div_exp_phase’, ’num_div_stat_phase’, ’Time_of_last_division’. Data augmentation was performed by successively (1) calculating all possible log2 ratios between features for each frame, (2) calculating the log2 fold changes for each initial feature between each pair of consecutive frames and (3) calculating time window (5 frames *i.e.* 50 min) metrics (average, minimum, maximum) for all these log2 scaled features. Given the sensitivity of plsr to class imbalance, several small predictive model were constructed with equal numbers of persister (n=206) and sensitive (n=206) cell lineages. To explore whether persisters resemble sensitive cells that die early or late in the post-treatment phase, multiple subsets of sensitive cells were sampled based on incremental time-of-death windows. Plsr analyses and feature reduction were performed as described above. The score of model contribution for each feature consists in the percentage of feature reduction cycles where the feature has been kept (100% means that the feature has been retained in all feature reduction models) (Figure S1c).

## Ethics statement

The clinical *P. aeruginosa* isolates used in this study were cultured from patient samples collected for routine microbiological testing at the University Hospital, Basel, Switzerland. Sub culturing and analysis of bacteria were performed anonymously. No additional procedures were carried out on patients. Cultures were sampled by following regular procedures with written informed consent, in agreement with the guidelines of the Ethikkommission beider Basel EKBB.

## Data availability

Mass spectrometry proteomic .raw files and accompanying metadata can be accessed at https://massive.ucsd.edu/ under the accession MSV000091240.

## Acknowledgments

Authors are thankful to Sebastian Hiller and Raphael Böhm for valuable help and providing unpublished results; Michelle Roulier, Martina Bläsi and Fabienne Hamburger for their technical support and Alexander Klotz for its expertise in microbial genetics. This study was supported by the Swiss National Science Foundation NRP72 project grant (407240_167080 to U.J.) and by the Swiss National Science Foundation NCCR AntiResist (51NF40_180541 to U.J.). We declare no competing financial interests.

