## Supplementary figures S1-S7 for "Coenzyme A depletion causes antibiotic tolerance in *Pseudomonas aeruginosa*"

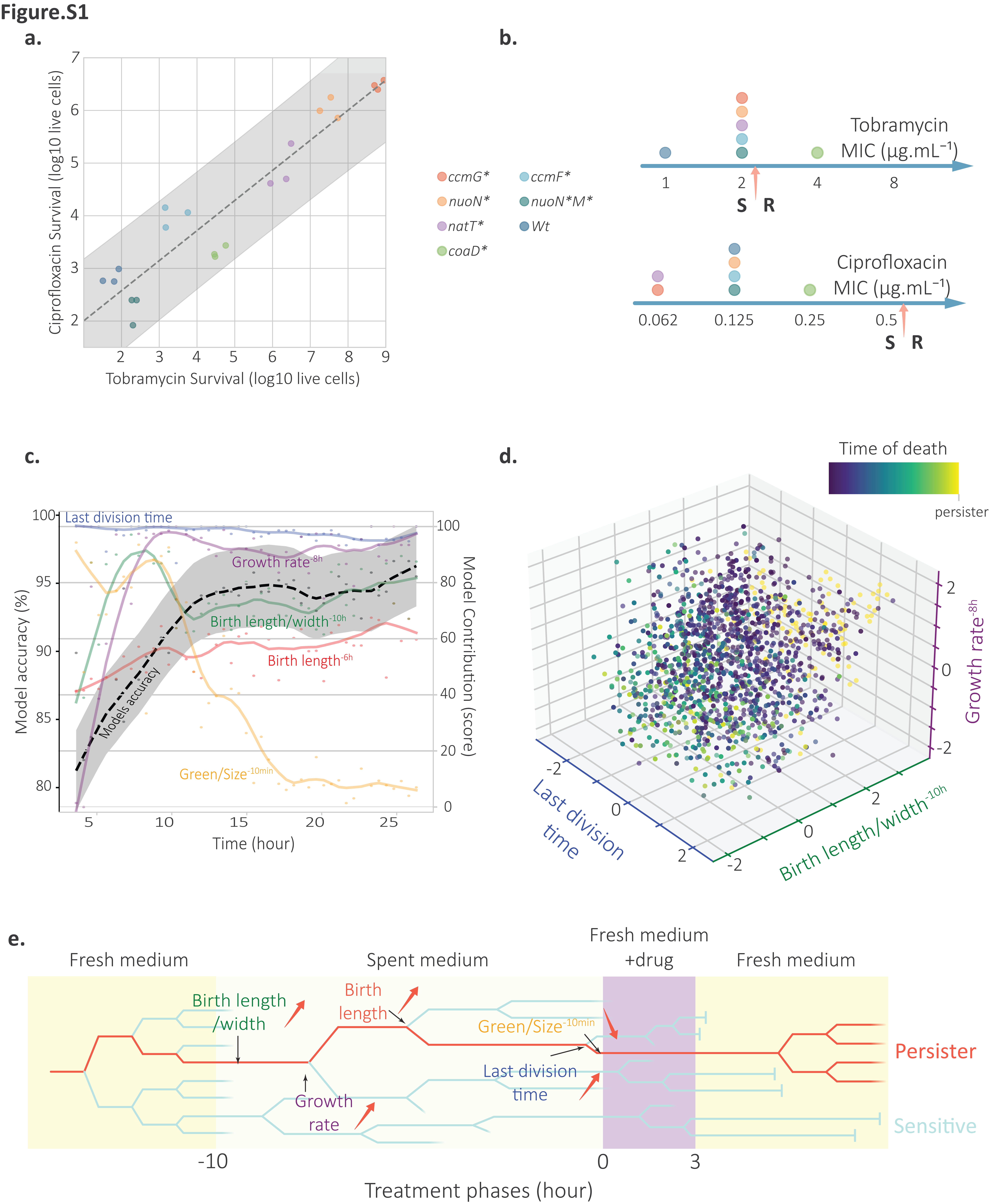

**Figure S1: Specific features of hyper-tolerant strains of *P. aeruginosa*. (a)** Hyper-tolerant strains of *P. aeruginosa* obtained though experimental evolution show multidrug tolerance. Survival of individual strains during treatment with tobramycin (X-axis) and ciprofloxacin (Y-axis) (Log10) is plotted for three independent replicas. **(b)** Minimal inhibitory concentration of hyper-tolerant strains from panel (a). Arrows indicate resistance Breakpoints as set by the EUCAST (2022). **(c)** Models predicting *natT** persisters based on microscopic features recorded prior treatment for sensitive cells disintegrating at different time after treatment. Because plsr requires data sets with equilibrated classes, each model was independently computed for different data subsets composed of 206 persister and 206 sensitive (out of 8423). Data slices were constructed with increasing sensitive-cell disruption time after treatment window (X-axis). Models accuracy for the different datasets were plotted in black (dotted curve, LOESS regression) (left Y-axis). Colored dots and curves (LOESS regression) indicate the contribution of specific discriminating microscopic features (see methods) for the prediction of survival/killing within specific time windows of sensitive-cell disruption time (right Y-axis). **(d)** 3D XYZ plot of individual lineages (dots) of the *natT** mutant strain recorded in microfluidic experiments (see Figure 1d,e) based on the three best persister-discriminating features in plsr models. These include the normalized ratio between the cell size at birth divided by the width 10 h prior treatment (X-axis), the normalized growth rate 8 h prior treatment (Y-axis), and the normalized time of last division prior treatment (higher = closer to treatment) (Z-axis). The sensitive-cell disruption time of individual lineages is indicated in color with yellow cells representing persisters (*i.e.* that never disrupted). **(e)** Schematic of a persister lineage during different phases of growth and treatment in microfluidic devices as indicated in Figure 1d,e. Red arrows indicate direction of variation for corresponding features in persister lineages: increased (up) or decreased (down).

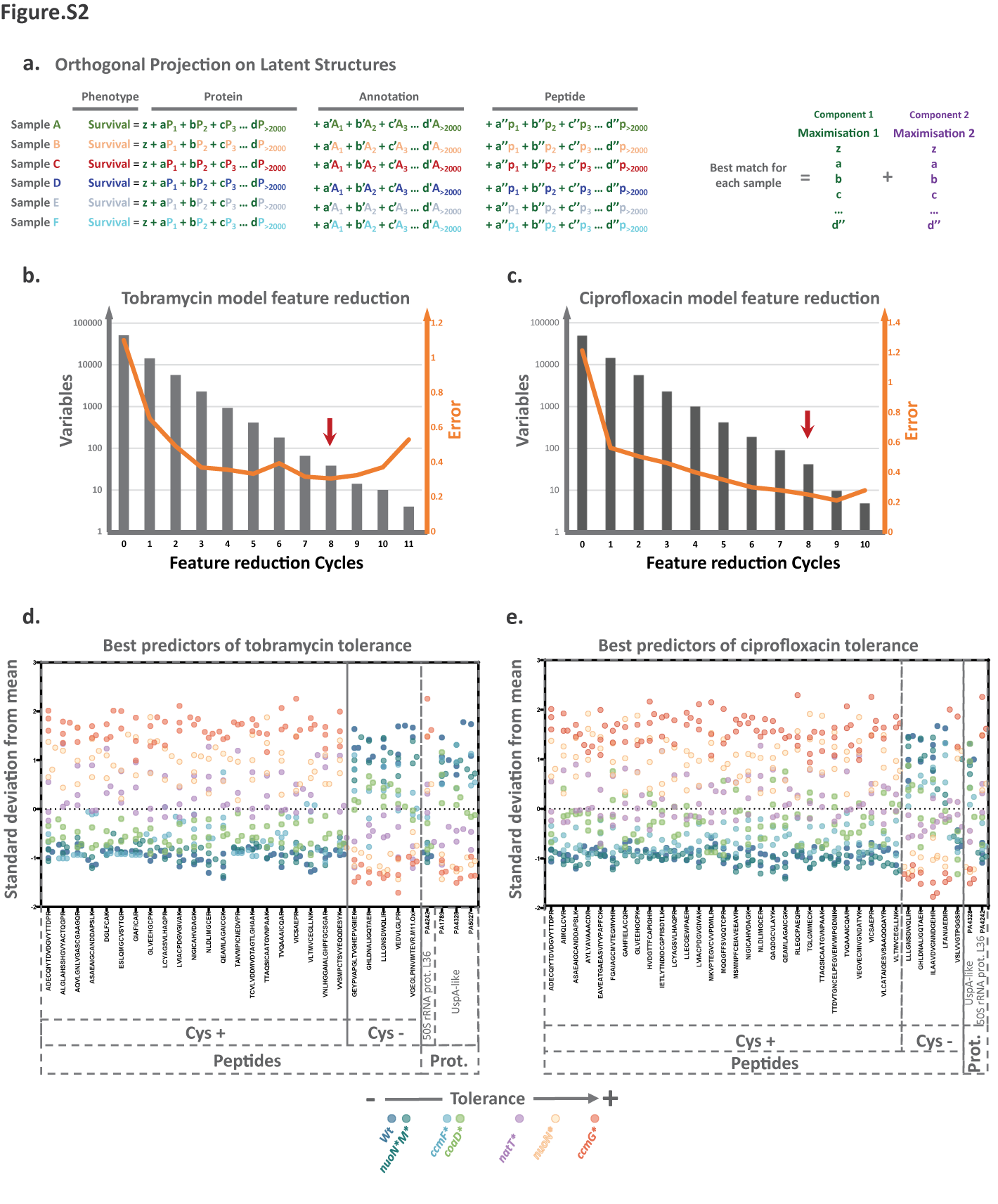

**Figure S2: Theoretical background of the Projection on Latent Structures Regression (plsr) model. (a)** Projection on Latent Structures Regression. The phenotypes (survival to antibiotic exposure) of different strains (A-F) are modeled as a combination of linear functions of MS intensities of peptides or proteins (sum of peptides from a given protein) and annotations (sum of peptides from proteins sharing the same annotation) (see material and methods). The plsr algorithm successively extracts components (*i.e.* linear functions) that maximize the covariance between omics variables and the predicted phenotype for the first component and residual unexplained phenotype values for the following components. Number of components were limited to two to avoid overfitting and facilitate biological interpretation of the models. Omics variables with the lowest model contribution (quantified by Variable Importance in Projection (VIP)) were excluded iteratively to compute simpler models with a smaller subset of variables. **(b-c)** Cycles of feature reduction for tobramycin **(b)** and ciprofloxacin **(c)** tolerance models. Grey bars (left Y-axis) represent numbers of remaining variables for each cycle of variable reduction (X-axis). Orange lines (right Y-axis) denote the quality of prediction (root mean square error of prediction from leave-one-out cross validations) with the selected model marked by a red arrow (see Figure 2). **(d-e)** Plotting of the most predictive biomarkers for survival during tobramycin **(d)** or ciprofloxacin **(e)** treatment. Colored dots represent single replicates (n=3) of protein mass spectrometry intensities (Y-axis) for 7 different hyper-tolerant strains as indicated in the legend (bottom). Cys+: peptides with at least one cysteine; Cys-: peptides without cysteines; Full-length proteins (Prot.) used by the models are indicated.

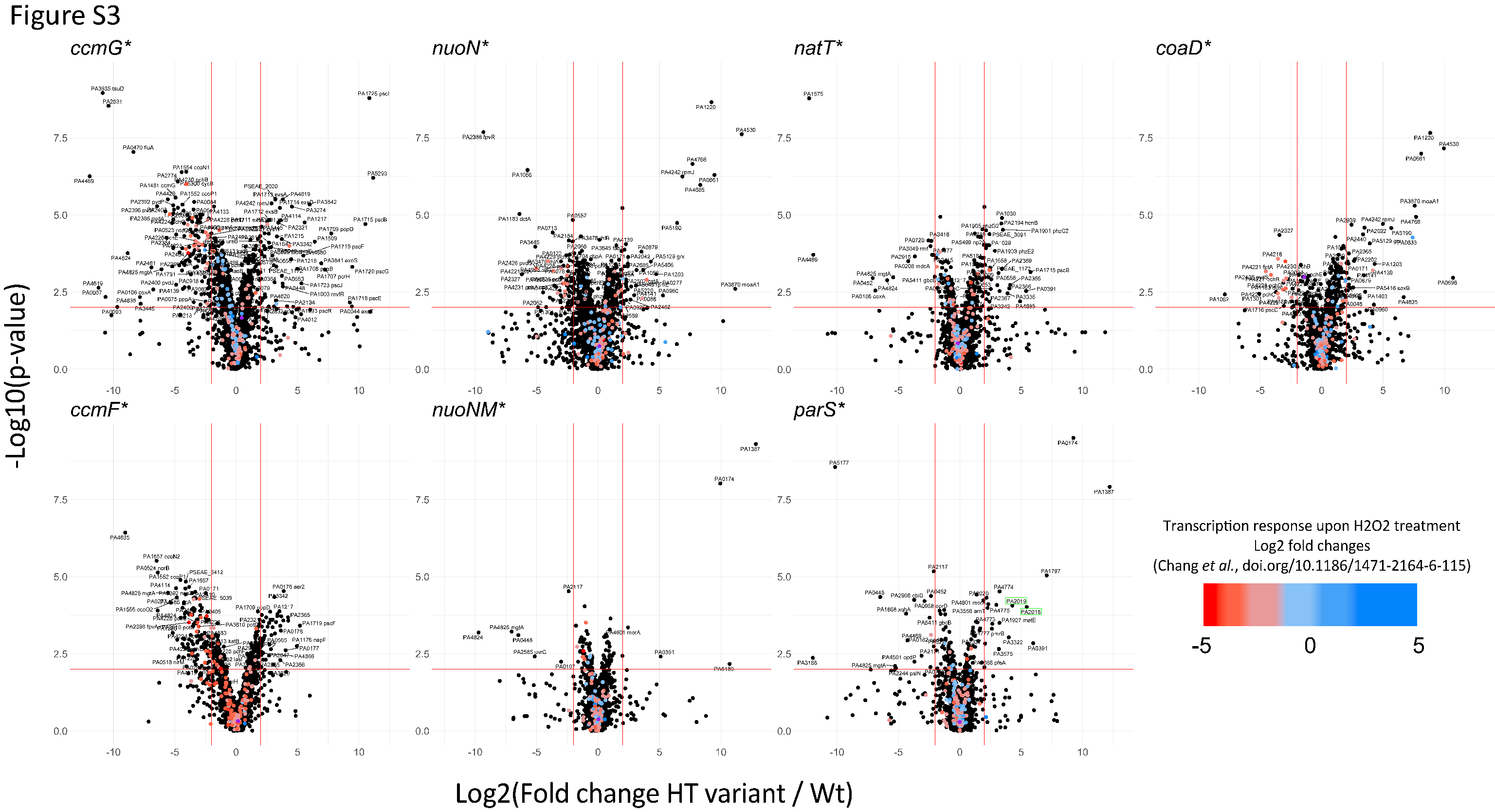

**Figure S3: The proteomic redox stress response of hyper-tolerant variants is highly variable.** Proteomic profiles of hyper-tolerant *P. aeruginosa* strains presented in this study ([massive.ucsd.edu:](https://massive.ucsd.edu/) MSV000091240). Dots indicate mass spectrometry intensity fold changes of proteins from the indicated mutant as compared to *P. aeruginosa* wild type (Log2; X-axis) and associated significance values (-Log_10_(p-value); Y-axis). Average fold changes and p-values are based on 3 biological replicates. The gradient color scale and dots in individual plots indicate transcriptional changes of specific genes of *P. aeruginosa* upon oxidative stress as reported previously ^1^(fold changes > 2). CoaD (purple dot) and over-expressed Mex efflux pumps components (green boxes) are marked.

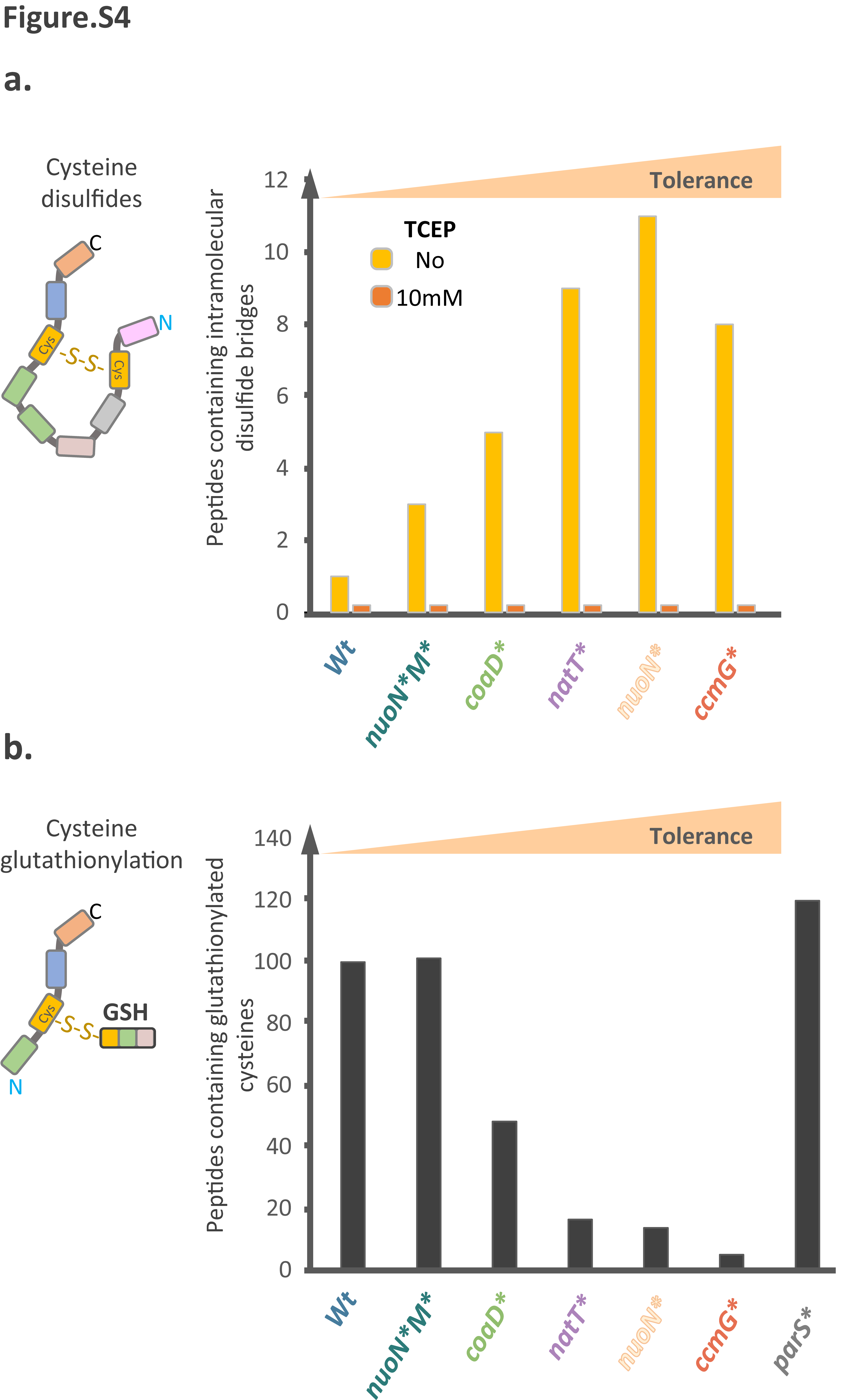

**Figure S4: Post-translational modifications of cysteine-containing peptides in different hyper-tolerant strains of *P. aeruginosa*. (a)** Correlation between antibiotic tolerance of different hyper-tolerant strains and peptides carrying an intramolecular cysteine disulfide bridge. Proteomic mass spectrometry identified peptides containing two cysteines and a mass shift corresponding to a disulfide bridge formation (here, -116 Da = two missing iodoacetamide carboxyamidomethylation + two missing protons) in the absence (yellow) but not in the presence (orange) of the reducing agent TCEP (Y-axis). The total number of observations was from pooled replicates (n=3). **(b)** Peptides carrying glutathionylated cysteine residues (determined by mass spectrometry) anticorrelate with overall tolerance levels.

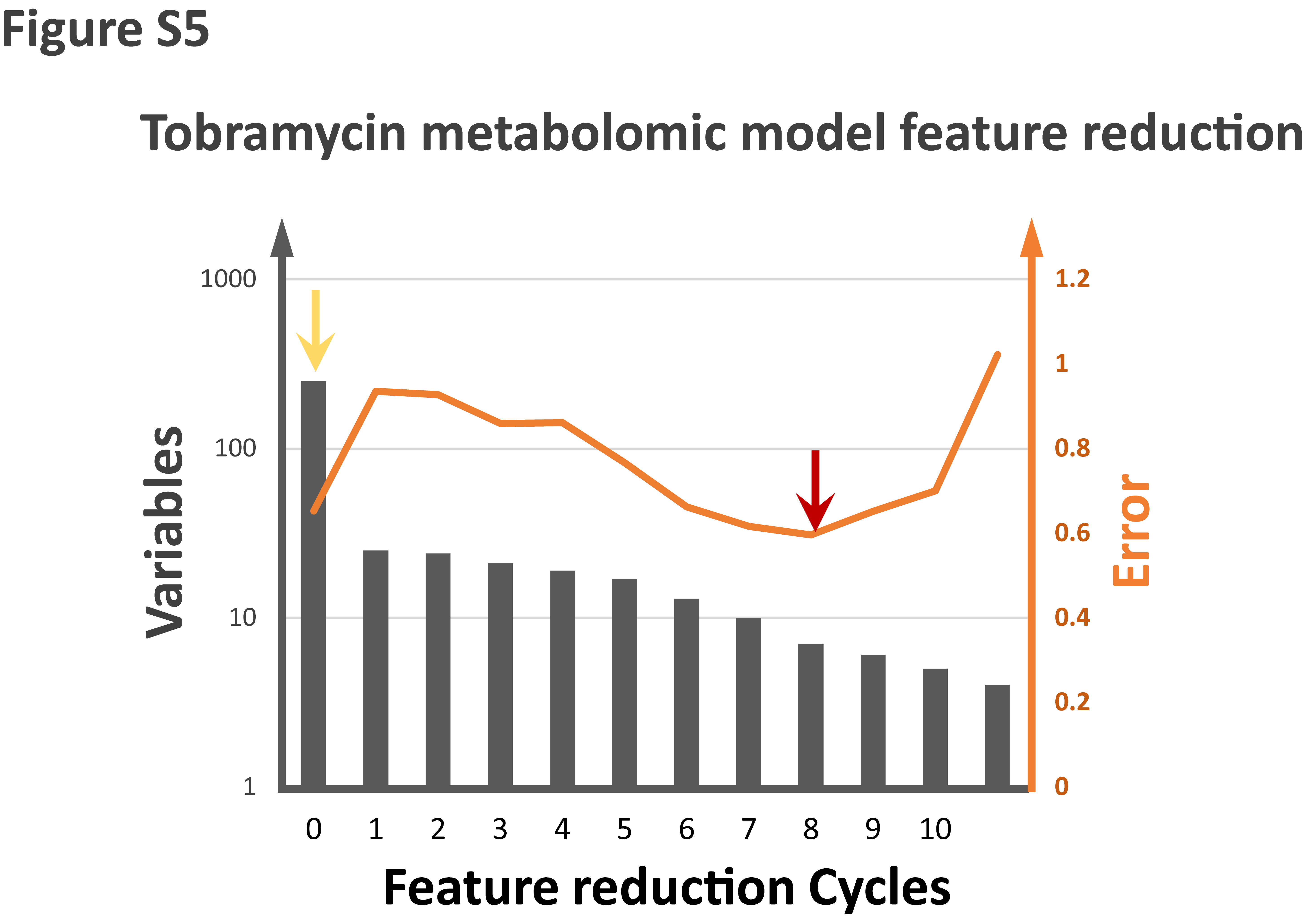

**Figure S5: Cycles of feature reduction for the metabolomics plsr model of tobramycin tolerance.** The number of variables in each model is represented by grey bars (left Y-axis) for each cycle of variable reduction (X-axis). Prediction quality (root mean square error) is indicated by the orange line (right Y-axis). Initial model (yellow arrow) was computed with 252 annotated metabolites and corresponds to Figure 3b. The best predictive model (red arrow) corresponds to the cross validation in Figure 3a.

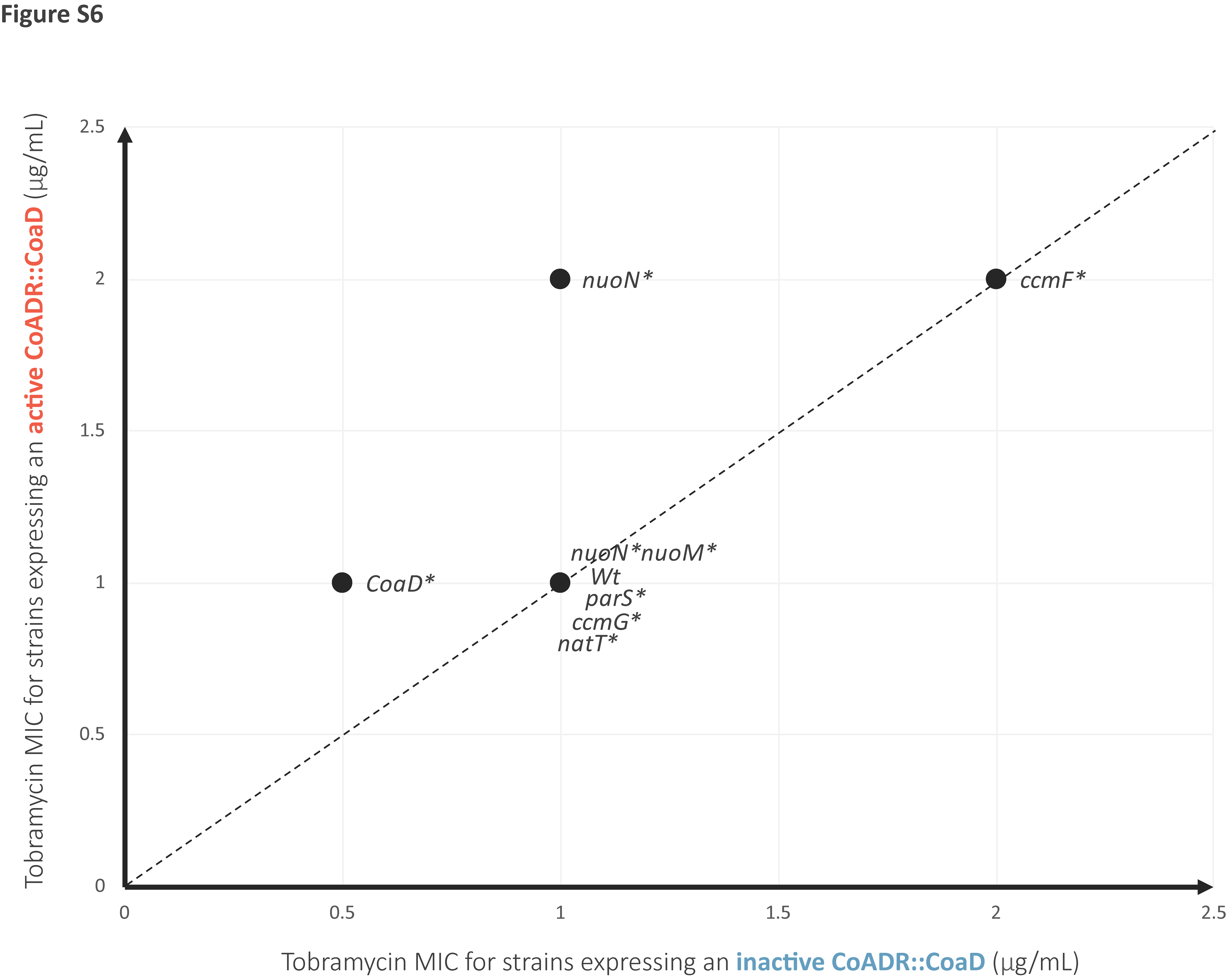

**Figure S6: MIC of tobramycin for *P. aeruginosa* strains expressing the active or the inactive version of the CoADR::CoaD construct.** Minimum inhibitory concentration for tobramycin (in μg/mL) in LB medium containing tetracycline at 100 μg/mL and 1mM IPTG to ensure expression of the active (Y-axis) or inactive (X-axis) fusion construct of CoADR::CoaD.

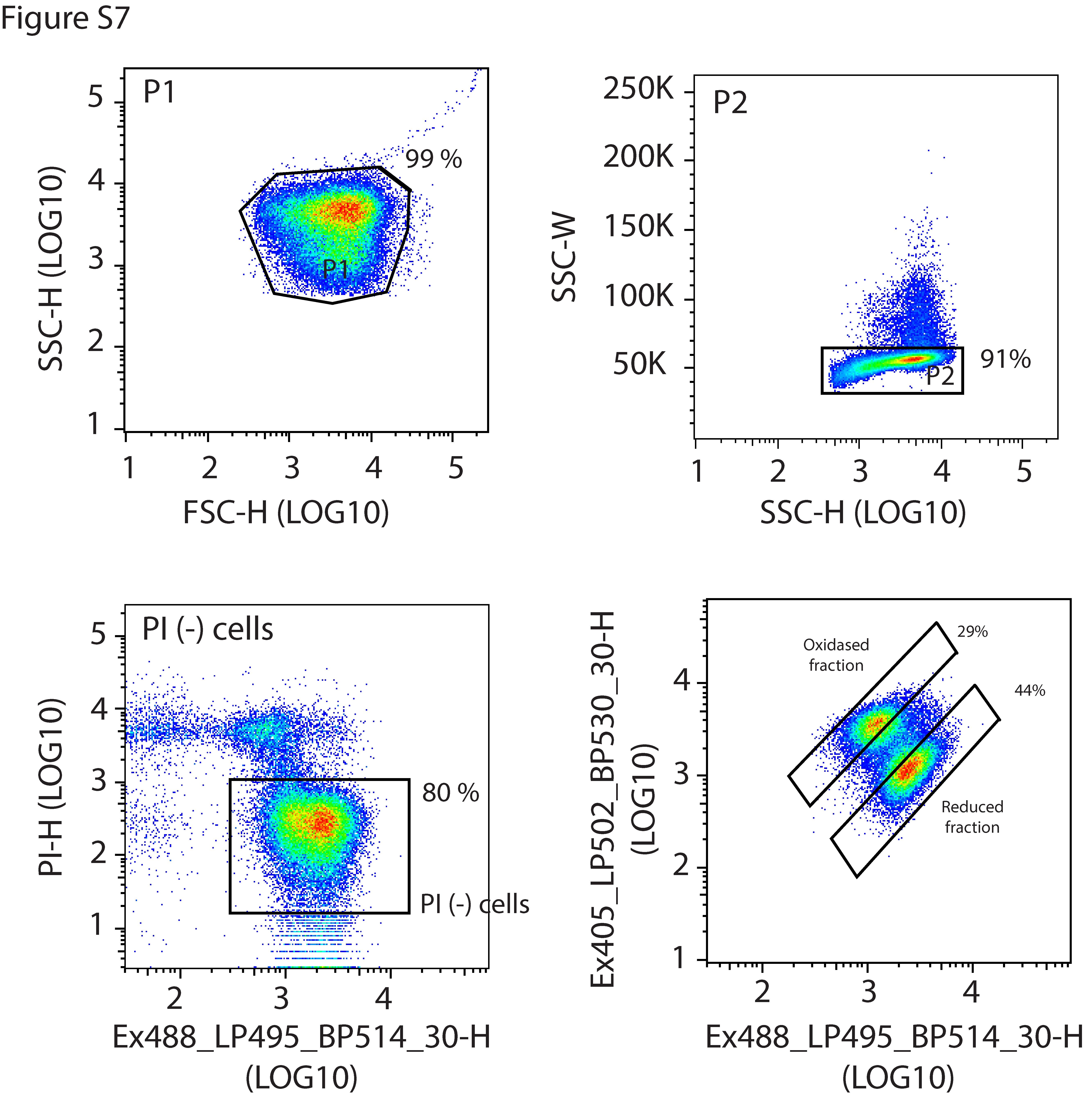

**Figure S7: Gating strategy of *P. aeruginosa* cells with different redox states.** FSC and SSC gating strategy for single bacteria capture (P1 and P2) is shown. Viable cells (propidium iodide negative; PI (-)) and GFP-positive cells were captured on gates defined with the help of non-fluorescent samples. Fraction of total populations captured are indicated. Gates for oxidized and reduced populations capture the large majority of fluorescent cells at different signals ratio between the Ex405_LP502_BP530/30-H and the Ex488_LP495_BP514/30-H channels.
