## Supplementary material for "Coenzyme A depletion causes antibiotic tolerance in *Pseudomonas aeruginosa*": Table S4

### Supplementary Table 4. List of strains and plasmids used in this study

| **Strains** | **Description** | **Reference** |
| --- | --- | --- |
| ***Pseudomonas aeruginosa*** | | |
| PAO1 | Wilde type *P. aeruginosa* | (Holloway 1955) |
| *nuoN** | PAO1 variant with a G300A mutation in NuoN protein | (Santi et al. 2021) |
| *nuoN*M** | PAO1 variant *nuoN** with a deletion of F184 in NuoM protein | (Santi et al. 2021) |
| *parS** | PAO1 variant with a G388D in the ParS protein | (Santi et al. 2021) |
| *ccmG** | PAO1 variant with a stop codon at residue 49 in the CcmG protein | (Santi et al. 2021) |
| *coaD** | PAO1 variant with a T71Q mutation in the CoaD protein | (Santi et al. 2021) |
| *ccmF** | PAO1 variant with a 403MAAL407 deletion in the PA1480 protein | (Santi et al. 2021) |
| *natT** | PAO1 variant with a E29D mutation in the PA1030 protein | (Santi et al. 2021) |
| PAO1*-roGFP2* | PAO1 transformed with pME6032::*Px2::roGFP2* | This study |
| *nuoN*M***-roGFP2* | *nuoN*M** transformed with pME6032::*Px2::roGFP2* | This study |
| *coaD*-roGFP2* | *coaD** transformed with pME6032::*Px2::roGFP2* | This study |
| *natT*-roGFP2* | *natT** transformed with pME6032::*Px2::roGFP2* | This study |
| PAO1-KD-*coaD* | Wilde type *P. aeruginosa* PAO1 with a chromosomal *Plac*::*dCas9* insert in the glmS1 neutral locus transformed with pBx-Spas-*sgRNA.PA0363.2* | This study |
| PAO1-KD-*mock* | Wilde type *P. aeruginosa* PAO1 with a chromosomal *Plac*::*dCas9* insert in the glmS1 neutral locus transformed with pBx-Spas-*sgRNA* (*i.e.* no sgRNA) | This study |
| *natT*-VC* | *natT** transformed with the “empty” intact pME6032 | This study |
| *natT*-gfp* | *natT** transformed with pME6032::*gfp* | This study |
| *natT*-coaD_pae_* | *natT** transformed with pME6032::*coaD_pae_* | This study |
| *natT*-coaD_st_* | *natT** transformed with pME6032::*coaD_st_* | This study |
| *natT*-coADR_st_* | *natT** transformed with pME6032::*coADR_st_* | This study |
| *natT*-coADR-D_act_* | *natT** transformed with pME6032::*coADR-D_act_* | This study |
| *natT*-coADR-D_ina_* | *natT** transformed with pME6032::*coADR-D_ina_* | This study |
| *ccmF*-coADR-D_act_* | *ccmF** transformed with pME6032::*coADR-D_act_* | This study |
| *ccmF*-coADR-D_ina_* | *ccmF** transformed with pME6032::*coADR-D_ina_* | This study |
| *coaD*-coADR-D_act_* | *coaD** transformed with pME6032::*coADR-D_act_* | This study |
| *coaD*-coADR-D_ina_* | *coaD** transformed with pME6032::*coADR-D_ina_* | This study |
| *nuoN*-coADR-D_act_* | *nuoN** transformed with pME6032::*coADR-D_act_* | This study |
| *nuoN *-coADR-D_ina_* | *nuoN** transformed with pME6032::*coADR-D_ina_* | This study |
| *nuoNM*-coADR-D_act_* | *nuoNM** transformed with pME6032::*coADR-D_act_* | This study |
| *nuoNM*-coADR-D_ina_* | *nuoNM** transformed with pME6032::*coADR-D_ina_* | This study |
| *parS*-coADR-D_act_* | *parS** transformed with pME6032::*coADR-D_act_* | This study |
| *parS*-coADR-D_ina_* | *parS** transformed with pME6032::*coADR-D_ina_* | This study |
| *Clin.1* | Clinical strain isolated from a CF patient sputum | This study |
| *Clin.5* | Clinical strain isolated from a CF patient sputum | This study |
| *Clin.15* | Clinical strain isolated from a CF patient sputum | This study |
| *Clin.22* | Clinical strain isolated from a CF patient sputum | This study |
| *Clin.24* | Clinical strain isolated from a CF patient sputum | This study |
| *Clin.27* | Clinical strain isolated from a CF patient sputum | This study |
| *Clin.28* | Clinical strain isolated from a CF patient sputum | This study |
| *Clin.52* | Clinical strain isolated from a CF patient sputum | This study |
| *Clin.54.A* | Clinical strain isolated from a CF patient sputum | This study |
| *Clin.54.B* | Clinical strain isolated from a CF patient sputum | This study |
| *Clin.58* | Clinical strain isolated from a CF patient sputum | This study |
| *Clin.61* | Clinical strain isolated from a CF patient sputum | This study |
| *Clin.71* | Clinical strain isolated from a CF patient sputum | This study |
| *Clin.75* | Clinical strain isolated from a CF patient sputum | This study |
| *Clin.83.A* | Clinical strain isolated from a CF patient sputum | This study |
| *Clin.83.B* | Clinical strain isolated from a CF patient sputum | This study |
| *Clin.130* | Clinical strain isolated from a CF patient sputum | This study |
| *Clin.131* | Clinical strain isolated from a CF patient sputum | This study |
| *Clin.132* | Clinical strain isolated from a CF patient sputum | This study |
| *Clin.182* | Clinical strain isolated from a CF patient sputum | This study |
| *Clin.195* | Clinical strain isolated from a CF patient sputum | This study |
| *Clin.218* | Clinical strain isolated from a CF patient sputum | This study |
| *Clin.225* | Clinical strain isolated from a CF patient sputum | This study |
| *Clin.226* | Clinical strain isolated from a CF patient sputum | This study |
| *Clin.234* | Clinical strain isolated from a CF patient sputum | This study |
| *Clin.239* | Clinical strain isolated from a CF patient sputum | This study |
| *Clin.263* | Clinical strain isolated from a CF patient sputum | This study |
| *Clin.265* | Clinical strain isolated from a CF patient sputum | This study |
| *Clin.266* | Clinical strain isolated from a CF patient sputum | This study |
| *Clin.277* | Clinical strain isolated from a CF patient sputum | This study |
| *Clin.327* | Clinical strain isolated from a CF patient sputum | This study |
| *Clin.338* | Clinical strain isolated from a CF patient sputum | This study |
| *Clin.342.A* | Clinical strain isolated from a CF patient sputum | This study |
| *Clin.342.B* | Clinical strain isolated from a CF patient sputum | This study |
| *Clin.344* | Clinical strain isolated from a CF patient sputum | This study |
| *Clin.351* | Clinical strain isolated from a CF patient sputum | This study |
| *Clin.483* | Clinical strain isolated from a CF patient sputum | This study |
| *Clin.531* | Clinical strain isolated from a CF patient sputum | This study |
| *Clin.649* | Clinical strain isolated from a CF patient sputum | This study |
| *Clin.654* | Clinical strain isolated from a CF patient sputum | This study |
| *Clin.686.A* | Clinical strain isolated from a CF patient sputum | This study |
| *Clin.686.B* | Clinical strain isolated from a CF patient sputum | This study |
| *Clin.687* | Clinical strain isolated from a CF patient sputum | This study |
| *Clin.694* | Clinical strain isolated from a CF patient sputum | This study |
| *Clin.695.A* | Clinical strain isolated from a CF patient sputum | This study |
| *Clin.695.B* | Clinical strain isolated from a CF patient sputum | This study |
| *Clin.695.C* | Clinical strain isolated from a CF patient sputum | This study |
| *Clin.699.A* | Clinical strain isolated from a CF patient sputum | This study |
| *Clin.699.B* | Clinical strain isolated from a CF patient sputum | This study |
| *Clin.706.A* | Clinical strain isolated from a CF patient sputum | This study |
| *Clin.706.B* | Clinical strain isolated from a CF patient sputum | This study |
| *Clin.710* | Clinical strain isolated from a CF patient sputum | This study |
| *Clin.725.A* | Clinical strain isolated from a CF patient sputum | This study |
| *Clin.725.B* | Clinical strain isolated from a CF patient sputum | This study |
| *Clin.725.C* | Clinical strain isolated from a CF patient sputum | This study |
| *Clin.731.A* | Clinical strain isolated from a CF patient sputum | This study |
| *Clin.731.B* | Clinical strain isolated from a CF patient sputum | This study |
| *Clin.732* | Clinical strain isolated from a CF patient sputum | This study |
| *Clin.755* | Clinical strain isolated from a CF patient sputum | This study |
| *Clin.766* | Clinical strain isolated from a CF patient sputum | This study |
| *Clin.775* | Clinical strain isolated from a CF patient sputum | This study |
| *Clin.781* | Clinical strain isolated from a CF patient sputum | This study |
| *Clin.812* | Clinical strain isolated from a CF patient sputum | This study |
| *Clin.1043* | Clinical strain isolated from a CF patient sputum | This study |
| *Clin.1047* | Clinical strain isolated from a CF patient sputum | This study |
| *Clin.1060* | Clinical strain isolated from a CF patient sputum | This study |
| *Clin.1124* | Clinical strain isolated from a CF patient sputum | This study |
| *Clin.1136* | Clinical strain isolated from a CF patient sputum | This study |
| *Clin.27-coADR-D_act_* | *Clin.27* transformed with pME6032:: *coADR-D_act_* | This study |
| *Clin.54.A-coADR-D_act_* | *Clin.54.A* transformed with pME6032:: *coADR-D_act_* | This study |
| *Clin.54.B-coADR-D_act_* | *Clin.54.B* transformed with pME6032:: *coADR-D_act_* | This study |
| *Clin.699-coADR-D_act_* | *Clin.699* transformed with pME6032:: *coADR-D_act_* | This study |
| *Clin.695.A-coADR-D_act_* | *Clin.695.A* transformed with pME6032:: *coADR-D_act_* | This study |
| *Clin.695.B-coADR-D_act_* | *Clin.695.B* transformed with pME6032:: *coADR-D_act_* | This study |
| *Clin.695.C-coADR-D_act_* | *Clin.695.C* transformed with pME6032:: *coADR-D_act_* | This study |
| *Clin.132.A-coADR-D_act_* | *Clin.132.A* transformed with pME6032:: *coADR-D_act_* | This study |
| *Clin.132.B-coADR-D_act_* | *Clin.132.B* transformed with pME6032:: *coADR-D_act_* | This study |
| *Clin.1136.A-coADR-D_act_* | *Clin.1136.A* transformed with pME6032:: *coADR-D_act_* | This study |
| *Clin.1136.B-coADR-D_act_* | *Clin.1136.B* transformed with pME6032:: *coADR-D_act_* | This study |
| *Clin.239-coADR-D_act_* | *Clin.239* transformed with pME6032:: *coADR-D_act_* | This study |
| *Clin.531-coADR-D_act_* | *Clin.531* transformed with pME6032:: *coADR-D_act_* | This study |
| *Clin.766-coADR-D_act_* | *Clin.766* transformed with pME6032:: *coADR-D_act_* | This study |
| ***Escherichia coli*** |  |  |
| Cloning and plasmid storage strain DH5α | *endA1*, *hsdR17*(rK‐mK+), *supE44*, *recA1*, *gyrA* (*Nalr*), *relA1*, *Δ*(*lacIZYA‐argF*)U169, *deoR*, *Φ80dlacΔ(lacZ)M15* | (Woodcock et al. 1989) |
| **Plasmids** |  |  |
| pME6032 | Tet^R^, P_K_, 9.8 kb pVS1 derived shuttle vector | (Heeb et al. 2000) |
| pME6032::*Px2::roGFP2* | pME6032 expressing a PAO1 codon adapted version of the fusion reporter Grx1::roGFP2 under *Px2* promotion with a PAO1-translation rate maximized synthetic RBS. | This study |
| pBx-Spas-*sgRNA* | CRISPRi expression vectors for constitutive expression of sgRNA devoid of sgRNA sequence. | (Tan, Reisch, and Prather 2018) |
| pBx-Spas-*sgRNA.PA0363.2* | CRISPRi expression vectors for constitutive expression of the sgRNA.PA0363.2 | This study |
| pME6032::*coaD_pae_* | pME6032 encoding the PAO1 *coaD* gene (PA0363) under *Plac* promotion and with a context specific maximized RBS. | This study |
| pME6032::*coaD_st_* | pME6032 expressing a PAO1 codon adapted version of CoaD from *S. aureus* NCTC 8325 under Plac promotion and with a context specific maximized RBS. | This study |
| pME6032::*coADR_st_* | pME6032 expressing a PAO1 codon adapted version of CoADR from *S. aureus* NCTC 8325 under Plac promotion and with a context specific maximized RBS. | This study |
| pME6032::*coADR-D_act_* | pME6032 expressing a PAO1 codon adapted fusion of the reductase domain from CoADR and the full length CoaD from *S. aureus* NCTC 8325 under Plac promotion and with a context specific maximized RBS. | This study |
| pME6032::*coADR-D_ina_* | pME6032 expressing a PAO1 codon adapted fusion of the reductase domain from CoADR and the full length CoaD from *S. aureus* NCTC 8325 carrying inactivating polymorphisms at active sites (See Table S2) under Plac promotion and with a context specific maximized RBS. | This study |
| pME6032*::gfp* | pME6032 expressing the monomeric green fluorescent protein mNeonGreen from PAO1 under Plac promotion. | This study |
